# PARP14 is an interferon (IFN)-induced host factor that promotes IFN production and affects the replication of multiple viruses

**DOI:** 10.1101/2024.04.26.591186

**Authors:** Srivatsan Parthasarathy, Pradtahna Saenjamsai, Hongping Hao, Anna Ferkul, Jessica J. Pfannenstiel, Daniel S. Bejan, Yating Chen, Ellen L. Suder, Nancy Schwarting, Masanori Aikawa, Elke Muhlberger, Adam J. Hume, Robin C. Orozco, Christopher S. Sullivan, Michael S. Cohen, David J. Davido, Anthony R. Fehr

**Affiliations:** Department of Molecular Biosciences, University of Kansas, Lawrence, Kansas 66045, USA; Department of Chemical Physiology and Biochemistry, Oregon Health Sciences University, Portland, OR, 97239, USA; Department of Molecular Biosciences, University of Texas, Austin, TX, 78712, USA; Department of Microbiology, Boston University School of Medicine, Boston, MA, 02118, USA; National Emerging Infectious Diseases Laboratories, Boston University, Boston, MA, 02118, USA; Center for Emerging Infectious Diseases Policy & Research, Boston University, Boston, MA, 02118, USA; Center for Excellence in Vascular Biology (P.K.J., M.A., E.A.), Brigham and Women’s Hospital, Harvard Medical School, Boston, MA, 02115, USA; Center for Interdisciplinary Cardiovascular Sciences (M.A., E.A.), Brigham and Women’s Hospital, Harvard Medical School, Boston, MA, 02115, USA; Channing Division of Network Medicine (M.A.), Brigham and Women’s Hospital, Harvard Medical School, Boston, MA, 02115, USA

**Keywords:** ADP-ribosylation, PARP, macrodomain, coronavirus, HSV-1, Ebola virus, Nipah virus, VSV, LCMV, interferon, interferon-stimulated genes

## Abstract

PARP14 is a 203 kDa multi-domain protein that is primarily known as an ADP-ribosyltransferase, and is involved in a variety of cellular functions including DNA damage, microglial activation, inflammation, and cancer progression. In addition, PARP14 is upregulated by interferon (IFN), indicating a role in the antiviral response. Furthermore, PARP14 has evolved under positive selection, again indicating that it is involved in host-pathogen conflict. We found that PARP14 is required for increased IFN-I production in response to coronavirus infection lacking ADP-ribosylhydrolase (ARH) activity and poly(I:C), however, whether it has direct antiviral function remains unclear. Here we demonstrate that the catalytic activity of PARP14 enhances IFN-*β* and IFN-*λ* responses and restricts ARH-deficient murine hepatitis virus (MHV) and severe acute respiratory syndrome coronavirus 2 (SARS-CoV-2) replication. To determine if PARP14’s antiviral functions extended beyond CoVs, we tested the ability of herpes simplex virus 1 (HSV-1), a DNA virus, vesicular stomatitis virus (VSV), a negative-sense RNA virus, and lymphocytic choriomeningitis virus (LCMV), an ambisense RNA virus, to infect A549 PARP14 knockout (KO) cells. While LCMV infection was unaffected, HSV-1 replication was increased in PARP14 KO cells and VSV replication was decreased. These results indicate that PARP14 restricts HSV-1 replication but enhances the replication of VSV. A PARP14 active site inhibitor had no impact on HSV-1 or VSV replication, indicating that its effect on these viruses was independent of its catalytic activity. These data demonstrate that PARP14 promotes IFN production and has both proviral and antiviral functions targeting multiple viruses.

**IMPORTANCE:** The antiviral response is largely regulated by post-translation modifications (PTM), including ADP-ribosylation. PARP14 is an ADP-ribosyltransferase that is upregulated by interferon and is under positive selection, indicating that it is involved in host-pathogen conflict. However, no anti-viral function has been described for PARP14. Here, we found that PARP14 represses both coronavirus and HSV-1 replication, demonstrating that PARP14 has antiviral functions. Surprisingly, we also found that PARP14 also has pro-viral functions, as it was critical for the efficient replication of VSV. These data indicate that PARP14 has both proviral and antiviral functions. Defining the mechanisms used by PARP14 to both repress and promote virus replication will provide new insights into how PARPs regulate virus infection.

## INTRODUCTION

Virus infection causes activation of host immune responses, with the interferon (IFN) response being the major pathway activated. Recognition of viral pathogen-associated molecular patterns (PAMPs) by host surface and cytosolic receptors leads to a series of molecular events that activates the expression of different types of IFNs, that eventually upregulates the expression of anti-viral IFN stimulated genes (ISGs). Several different post-translational modifications regulate this response, including phosphorylation, ubiquitination, and other less common modification such as ADP-ribosylation.

ADP-ribosylation is a post-translational modification (PTM) where ADP-ribose subunits are added to target proteins in a process catalyzed by ADP-ribosyltransferases (ARTs) that use NAD^+^ as a substrate. The ART Diphtheria toxin-like domain (ARTD) family of ART proteins are intracellular ARTs that are individually named PARPs [1]. These ARTDs catalyze the addition of either poly-ADP-ribose (PAR) or mono-ADP-ribose (MAR) to proteins or nucleic acid. The human genome encodes 17 PARPs that are involved in the regulation of most cellular activities including DNA damage repair, ER stress, cell cycle regulation, transcription and translation, and the innate and adaptive immune responses [1]. Apart from addition of ADP-ribose, removal of ADP-ribose from target proteins is also implicated in important cellular processes. The presence of ADP-ribose hydrolases (ARH) or macrodomains such as PARG, TARG1, MacroD1 and MacroD2 in humans underscores the importance of ADP-ribose turnover [2]. Furthermore, all coronaviruses (CoVs) encode for a highly conserved macrodomain (Mac1) that removes ADP-ribose from proteins, indicating that these viruses may be susceptible to antiviral ADP-ribosylation. Infection with murine hepatitis virus (MHV), SARS-CoV, or SARS-CoV-2 containing point mutants in key residues required for catalytic activity (i.e. MHV N1347A) or full deletions of Mac1 induce robust interferon responses and are highly attenuated *in vivo* [3-5]. In addition, the poor replication and enhanced induction of IFN following infection with MHV N1347A could be reversed using PARP inhibitiors [3]. These results demonstrated that PARP-mediated ADP-ribosylation is a potent anti-viral defense mechanism against CoVs.

The expression of several PARPs is activated by IFN in response to virus infection, categorizing them as interferon stimulated genes (ISGs). Some PARPs have known antiviral activity including PARP13 (zinc-antiviral protein or ZAP), PARP12, PARP11, PARP9, PARP5a/b, PARP7, and PARP1 [6]. In addition, some PARPs also act as pro-viral factors, promoting viral replication and infection. PARP11 was shown to suppress IFN-I signaling by MARylating IFNAR and priming it for ubiquitination and eventual degradation. The suppression of IFN-I signaling by PARP11 increased the replication of vesicular stomatitis virus (VSV) and herpes simplex virus 1 (HSV-1). These results demonstrated a pro-viral role for PARP11 [7]. PARP7-dependent ADP-ribosylation also suppressed IFN-I production and promoted influenza A virus infection, indicating that PARP7 also has pro-viral functions [8]. Consistent with this pro-viral function, PARP7 knockdown reduced the replication of murine hepatitis virus (MHV) A59, demonstrating its ability to promote coronavirus replication [3]. Finally, PARP1 inhibitors have also been shown to efficiently inhibit replication of several viruses such as herpesviruses, adenoviruses and human immunodeficiency virus (HIV), indicating that PARP1 may also promote virus replication in some cases [9-11].

PARP14 is an ISG that is important in several different biological processes including inflammation, cancer progression and DNA damage. PARP14 is a multidomain protein and its domain architecture was recently re-analyzed using Alphafold2 [12]. This analysis determined that PARP14 contains 3 RRM domains, 7 full KH domains and 1 split KH domain that are suspected to bind to nucleic acids, 3 macrodomains (MDs) one of which has an ADP-ribose hydrolase activity (MD1) [12-14], a WWE domain that is suspected to bind ADP-ribose subunits, and the ART catalytic domain [12].

PARP14 was originally identified as a STAT6-binding protein that promoted IL-4 dependent anti-inflammatory signaling [15]. Since then, several independent studies have implicated PARP14 in inflammatory and interferon signaling pathways. PARP14’s involvement in inflammatory and IFN responses is cell-type and insult dependent, suggesting that PARP14 has multiple functions that are context dependent. In M1 macrophages PARP14 suppresses IFNγ-induced activation of pro-inflammatory signaling by STAT1 MARylation [16]. We and others have found that PARP14 is also critical for inducing high levels of IFN-*β* following infection with *Salmonella typhimurium*, MHV, as well as treatments with LPS or poly(I:C) treatment [3, 17]. Specifically, PARP14 knockout M0 macrophages and epithelial cells demonstrated significantly reduced levels of IFN-*β* after infection or treatment with LPS or poly(I:C), indicating that one of PARP14’s functions is to promote IFN-*β* production, however the mechanisms by which it enhances IFN-*β* production remain unknown [3, 17]. More recently, several groups have demonstrated that PARP14 is the major PARP responsible for IFN-*γ* and poly(I:C) induced cytosolic ADP-ribose bodies (ICAB) [18-21].

Interestingly, PARP14 is one of the several PARPs that has evolved under positive selection, indicating that it may be involved in host-pathogen conflicts. This positive selection is seen in many of the different domains of PARP14, especially the macrodomains, WWE domain, and the catalytic domain at the C-terminus of the protein [22]. However, despite this evidence for involvement in host-pathogen interactions, there are very few reports describing any impact of PARP14 on virus or bacterial infections. In one report, PARP14 KO RAW264.7 macrophages had significantly increased levels of *S. typhimurium* replication [17]. In addition, siRNA knockdown of PARP14 led to a mild increase (∼2-fold) in the levels of a Mac1 mutant MHV viral RNA, but knockdown or a congenital knockout of PARP14 did not significantly increase the amount of infectious virus produced [3]. Despite PARP14 being a regulator of IFN production and triggers the production of ICABs, it remains unclear how or even if PARP14 impacts virus replication.

Recently, several tools have been developed to facilitate testing the impact of PARP14 on virus infection. First, we reported PARP14 knockout (KO) A549 and NHDF cell lines [3], which have intact IFN systems and are susceptible to infection by a wide variety of viruses. Second, a highly potent and selective PARP14 catalytic inhibitor, RBN01239, was developed [23]. Finally, we report here the creation of a tamoxifen-inducible PARP14 knockout mouse model. Hence, with the availability of an advanced PARP14 inhibitor and with the advent of several PARP14 knock-out systems, we sought to determine the effect of PARP14 and its catalytic activity on IFN production and the replication of multiple different virus families.

Utilizing these tools, we report here that PARP14 influences the replication of multiple viruses. PARP14 potently inhibited the replication of HSV-1 as well as Mac1-mutant MHV and SARS-CoV-2. Unexpectedly, PARP14 also acted as a proviral factor for vesicular stomatitis virus (VSV), a negative-sense RNA virus, highlighting the multifaceted effect of PARP14 on virus replication. Furthermore, we confirmed that the catalytic activity of PARP14 was critical for IFN-*β* and IFN-*λ* induction and the inhibition of CoV replication. This study highlights the involvement of PARP14 in the IFN response and virus infection, underscoring the importance of PARP14 as an immunomodulatory host factor that can affect the replication of multiple viruses.

## RESULTS

### PARP14 promotes IFN-*β* and IFN-*λ* production in poly(I:C) transfected cells

We previously demonstrated that PARP14 was required for IFN-*β* production in response to poly(I:C) in human A549 cells, via an indirect VSV bioassay [3]. To determine how PARP14 impacts IFN induction more quantitatively, we induced IFN production via poly(I:C) transfection and quantified the amount of IFN-β and IFN-*λ* transcripts by qPCR and protein by ELISA in A549 wild-type (WT) and PARP14 KO cells, that were developed using CRISPR by transducing A549 cells with control or PARP14 targeting guide RNAs and Cas9 [3]. Poly(I:C) transfection induced IFN-β **(Fig. 1A, S1A)** and IFN-λ **(Fig. 1B)** mRNA production in A549 cells, but their levels were significantly reduced in PARP14 KO compared to WT A549 cells. We also observed a similar decrease in IFN-β and IFN-λ protein levels following poly(I:C) transfection in PARP14 KO A549 cells compared to WT cells in an ELISA assay **(Fig. 1C-D)**. Poly(I:C) transfected PARP14 KO cells also had decreased protein levels of the interferon stimulated gene OAS3 compared to WT cells **(Fig. 1E-F)**. Finally, we also observed a nearly 4-fold reduction in IFN-*β* mRNA production in response to poly(I:C) transfection in PARP14 KO NHDF cells **(Fig. S1B)** [3]. Since PARP14 catalyzes MARylation, we used a highly potent and selective PARP14 catalytic inhibitor, RBN012759 (PARP14i) to test if the observed reduction in IFN and ISG levels were due to the MARylating activity of PARP14. PARP14i efficiently restricted PARP14’s auto-MARylation activity in PARP14 overexpressing 293T cells in a dose-dependent manner **(Fig. S1B)**. We utilized PARP14 G832E for these experiments as it was shown to be deficient in its MD1-dependent ADP-ribosyl hydrolase activity, which leads to enhanced autoMARylation [14]. Importantly, poly(I:C) stimulated A549 cells displayed a dose-dependent decrease in IFN-β transcript levels when treated with PARP14i. At the highest concentration of PARP14i, the level of IFN-β mirrored those in PARP14 KO cells **(Fig. 1G)**. This suggests that the reduction in IFN-β levels in PARP14 KO cells was due to the absence of PARP14 catalytic activity. In addition, we also used a PARP14 degrading compound, RBN012811 (DEG), and its associated negative control, RBN013527 (CTL), to test the role of PARP14 in IFN-*β* production (a generous gift from Mario Niepel, Ribon Therapeutics) [24]. DEG effectively reduced PARP14 protein to undetectable levels in WT A549 cells **(Fig. S1C)**. Compared to CTL, we found that the PARP14 DEG decreased IFN-*β* mRNA induction 3-fold, similar to results with complete PARP14 KO cells **(Fig. S1D)**. Finally, we restored IFN-β transcript and protein abundance to near WT levels by exogenously expressing full-length PARP14 in PARP14 KO cells **(Fig. 1H-I)**, further confirming that the decrease in IFN-β levels in PARP14 KO cells was not due to off-target effects of the PARP14 deletion. These results demonstrate that in response to poly(I:C) stimulation PARP14-dependent MARylation promotes IFN-*β* and IFN-*λ* production.

**Fig. 1.**
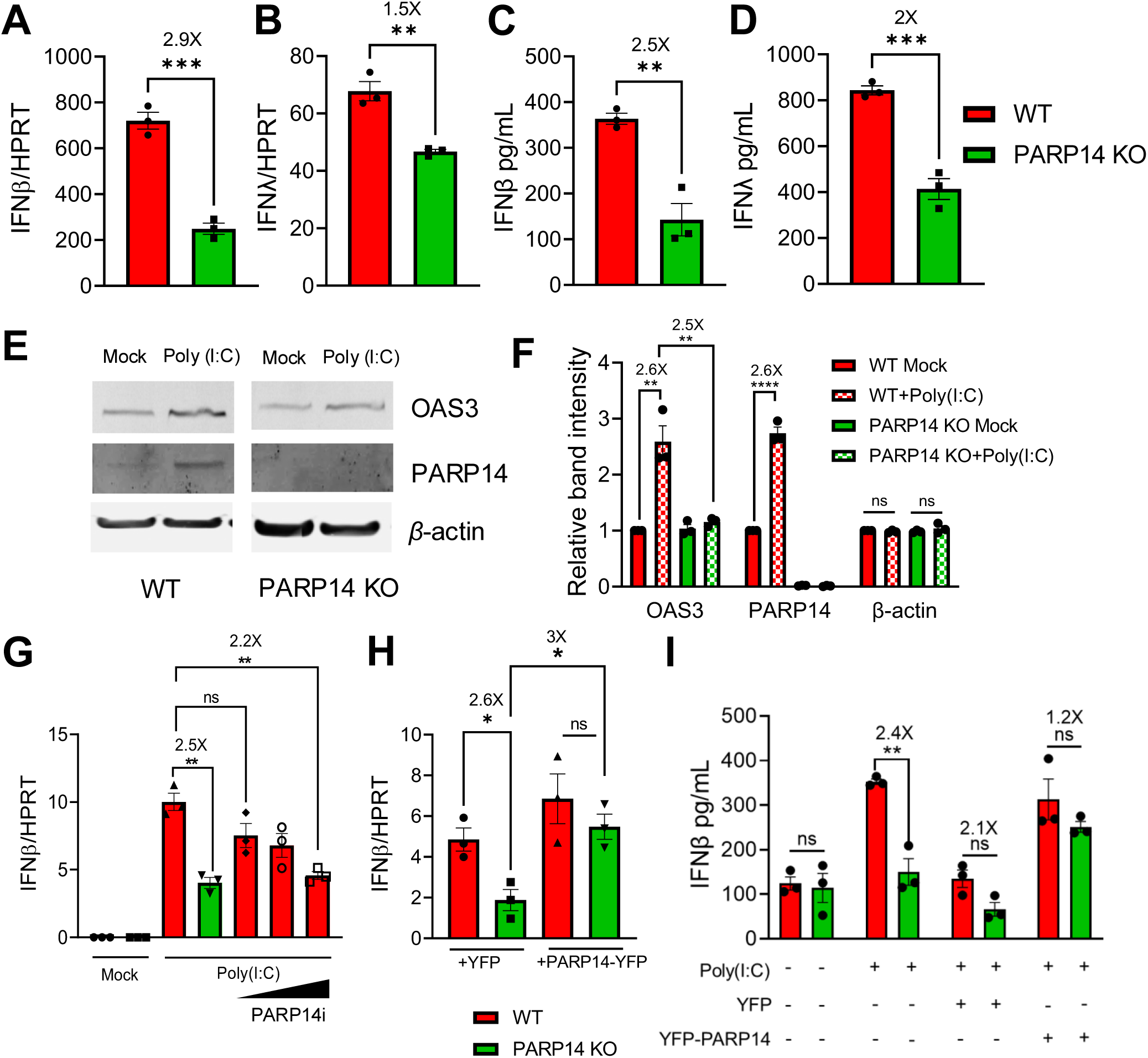
PARP14 promotes IFN production following poly(I:C) treatment in human A549 cells. A-B) WT (red bars) and PARP14 knock-out (KO) (green bars) A549 cells were transfected with 0.5ug/mL poly(I:C), RNA was isolated from cells at 18 hours post-tranfection (hpt) and IFNβ (A) and IFNλ (B) mRNA levels were quantified by qPCR using ΔCt method. C-D) WT and PARP14 KO cells were transfected with 0.5μg/mL poly(I:C). Supernatant was collected 18 hpt and IFNβ (C) and IFNλ (D) protein levels were quantified by ELISA. Data shown in A-D are from 1 experiment and are representative of 3 independent experiments with N=3 biological replicates for each experiment. E) WT and PARP14 KO cells were either mock transfected or transfected with 0.5 μg/mL of poly(I:C). Cell lysates were collected at 18 hpt and OAS3 and PARP14 protein levels were determined by immunoblotting using β-actin as a loading control. F) Band intensities from E were quantified by densitometry, normalized to β-actin, and then relative levels as compared to mock transfected WT cells were determined. Data shown in E-F are from 1 experiment and are representative of 3 independent experiments with N=1 biological replicate per group in each experiment. G) WT cells were either mock transfected or transfected with 0.5 μg/mL of poly(I:C) and were immediately either treated with DMSO (vehicle) or with increasing concentrations of PARP14i (0.01, 0.1, and 1 μM). At 18 hpt RNA was isolated from cells and IFNβ mRNA was quantified by qPCR using ΔCt method. H) WT A549 cells transfected with poly(I:C) were treated with 1 μM PARP14 Degron control (CTL) or PARP14 Degron (DEG) for 18 hours. RNA was isolated from cells at 18 hpt and IFNβ mRNA was quantified by qPCR using ΔCt method. I) WT and PARP14 KO cells were transfected with 0.5 μg of YFP or YFP-PARP14 plasmids. At 48 hpt cells were transfected with 0.5 μg/mL of poly(I:C) and 18 hours later RNA was isolated from cells and IFNβ mRNA was quantified by qPCR using ΔCt method. Data shown in G-I are from 1 experiment and are representative of 2-3 independent experiments with N=3 biological replicates per group for each experiment.

### PARP14 promotes IFN-*β* production in Mac1-mutant CoV infected cells

Using congenital PARP14 KO mouse cells, we previously showed that PARP14 was required to promote IFN-*β* production upon infection with MHV-N1347A [3]. To circumvent the limitations of constitutive PARP14 KO mice, which include poor breeding efficiency and potential confounding effects of developmental compensation for PARP14 KO by other PARPs, we developed a conditional tamoxifen-inducible PARP14 KO mouse model to better identify the role of PARP14 in response to MHV infection **(Fig. 2A)**. *LoxP* sites were inserted into the *parp14* locus around exon 2 via CRISPR mediated mutagenesis to create *parp14* floxed alleles. These mice were then bred with *ert2-cre* heterozygote mice to obtain both *parp14* floxed *ert2*-*cre* positive mice (*parp14^flox/Ert2-Cre^*), and as controls, *parp14* floxed ert2-*cre* negative offspring (*parp14^+/+^*) **(Fig. 2A)**. BMDMs were harvested from these mice and in *cre* positive cells induction with 4-hydroxy tamoxifen (4-OH-T) led to an efficient knock-out of PARP14 *(Parp14^-/-^)* **(Fig S2A)**. BMDMs that were *cre* negative still expressed PARP14 following treatment with 4-OH-T (*parp14^+/+^*). MHV-N1347A infection induced robust IFN-β levels compared to WT in *parp14^+/+^*BMDMs as previously demonstrated **(Fig. 2B)** [3]. However, in *parp14^-/-^* BMDMs infected with N1347A, the level of IFN-β was reduced to *parp14^+/+^* BMDM levels infected with WT virus, suggesting that PARP14 was required for IFN-β production in response to MHV-N1347A infection, confirming our prior observation and validating our KO mice **(Fig 2B)**. Similarly, there was also a decrease in the induction of ISG15 transcript levels in MHV-N1347A infected cells in *parp14^-/-^* cells **(Fig S2B)**. To determine if PARP14’s ART activity was responsible for the increase in IFN-*β* expression following infection, MHV-WT and N1347A infected BMDMs were treated with PARP14i and tested for IFN-β mRNA expression. PARP14i treatment caused a dose-dependent decrease in IFN-β expression in BMDMs infected with N1347A **(Fig. 2D)** similar to its impact on poly(I:C) induced IFNβ expression. Following MHV-N1347A infection, the level of secreted IFN-β protein was also reduced to WT levels in *Parp14^-/-^*BMDMs and in WT BMDMs treated with PARP14i **(Fig. 2E)**. Previously, we demonstrated that infection with a Mac1 deleted SARS-CoV-2 virus (ΔMac1) induced elevated levels of IFN-*β* and IFN-*λ* in human epithelial cells, which we hypothesized was due to PARP14 as it was highly upregulated following infection, though this was not formally proven [25]. Following ΔMac1 infection of Calu-3 cells, PARP14i treatment of Calu-3 cells reduced the mRNA levels of IFN-β, IFN-λ and CXCL-10 to levels seen following WT virus infection **(Fig. 2F)**. These data demonstrate that the increase in IFN-β levels in response to Mac1-deficient human and mouse CoV infection occurs via PARP14-dependent MARylation.

**Fig. 2.**
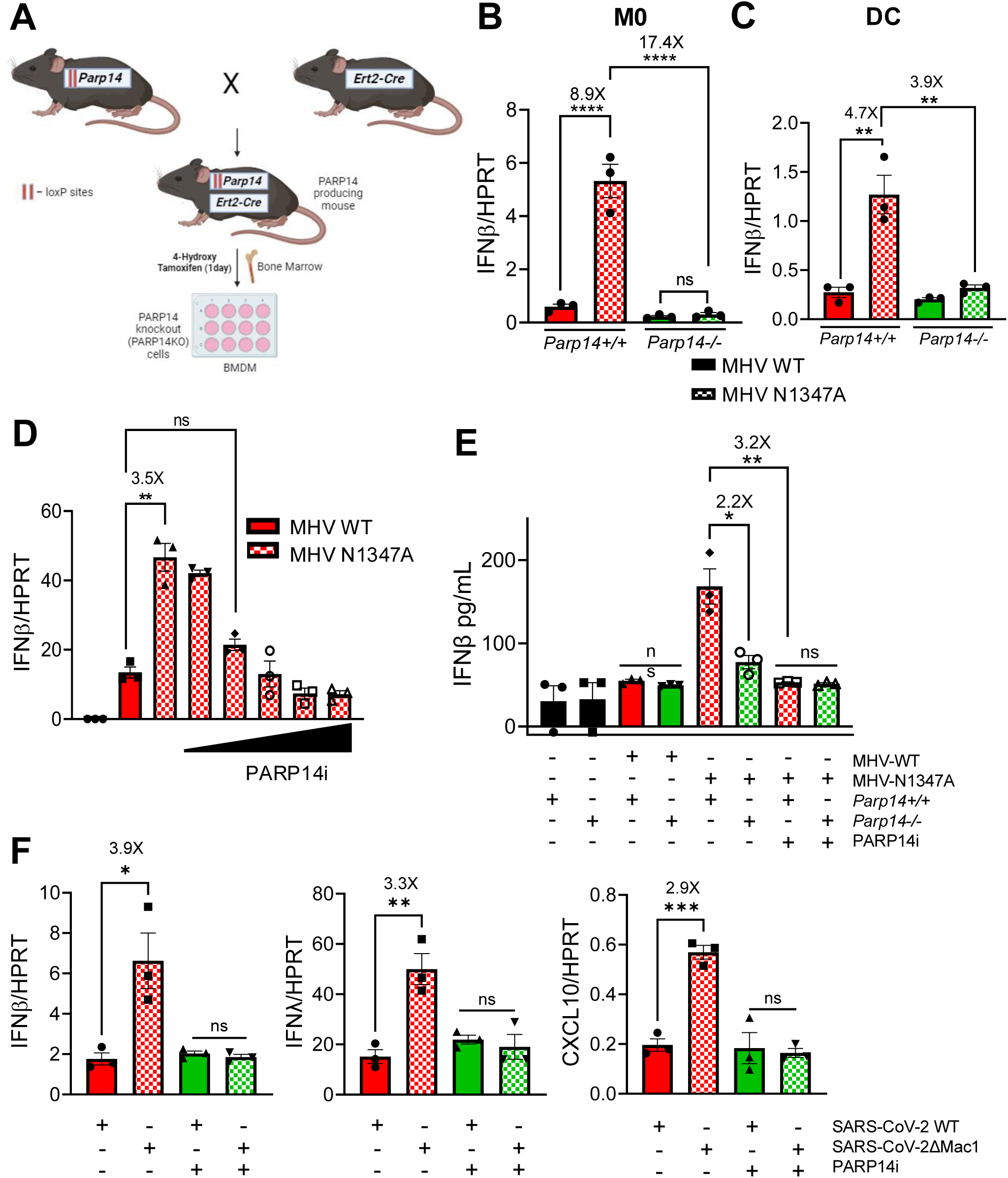
PARP14 promotes IFN production following Mac1 mutant CoV infection. A) Schematic diagram showing the generation of PARP14 KO mice for Bone-marrow cell isolation. *Parp14* floxed mice were crossed with *cre* heterozygote mice and the resulting progeny were used to harvest bone marrow. Bone marrow cells were isolated and plated with 10ng/mL of mCSF or GM-CSF for 6 days to differentiate cells into M0 macrophages or dendritic cells (DCs), respectively. On day 6, these cells were treated with 1μg/mL of 4-hydroxy tamoxifen (4-OHT) for 24 hours to induce *Cre* expression leading to the removal of exon 2 of *Parp14* (for details see Methods). B-C) *Parp14+/+* (Cre-) (red bars) and *Parp14-/-* (Cre+) (green bars) M0 cells (B) or DCs (C) were infected with MHV WT (solid bars) and N1347A (checkered bars) at an MOI of 0.1. RNA was isolated from cells at 12 hpi and IFNβ mRNA was quantified by qPCR using ΔCt method. D) *Parp14+/+* cells were infected with WT and N1347A at an MOI of 0.1 and then treated at 1 hpi with DMSO or increasing concentrations of PARP14 inhibitor (PARP14i) (3, 11, 33, 99 and 297nM). RNA was isolated from cells at 12 hpi and IFNβ mRNA levels were quantified using qPCR using ΔCt method. E) *Parp14+/+* and *Parp14-/-* M0 macrophages were infected with WT and N1347A MHV at an MOI of 0.1 and then treated with DMSO or 100 nM PARP14i at 1 hpi. The cell supernatant was collected at 12 hpi and IFNβ protein was quantified by ELISA. F) A549-ACE2 cells were infected with WT and *Δ*Mac1 SARS-COV-2 at an MOI of 0.1 and treated with DMSO or PARP14i. RNA was isolated from cells at 48 hpi and IFNβ, IFNλ and CXCL10 mRNA levels were quantified by qPCR using ΔCt method. Data shown in B-F are from 1 experiment and are representative of 3 independent experiments with N=3 biological replicates per group for each experiment.

### PARP14 promotes MDA-5 but not RIG-I dependent IFN production

IFN stimulation in virus-infected cells occurs due to recognition of viral RNA by pattern recognition receptors such as MDA5 or RIG-I. This leads to a cascade of events, resulting in activation of cytoplasmic proteins such as MAVS, TBK-1 and IRF3 and the nuclear translocation of IRF3 that leads to transcriptional activation of IFN production. BMDMs infected with N1347A MHV-JHM induced a robust production of IFN-β compared to WT infection. However, this difference in IFN-β production was lost during N1347A infection in BMDMs from MAVS knock-out (KO) (MAVS^-/-^) mice demonstrating the key role of intracellular RNA sensing for the induction of IFN-β **(Fig. 3A)**. We then tested the effect of MDA5 and RIG-I KO (Mda5^-/-^ and RIG-I^-/-^) on IFN-β production following infection with N1347A **(Fig. 3A)**. We found that RIG-I KO did not have any effect on IFN-β production during N1347A infection. In contrast, IFN-β levels were reduced to WT levels in Mda5^-/-^ BMDMs, suggesting the IFN-β production during N1347A infection occurred via the MDA5-dependent IFN-production pathway. Next, we tested to whether PARP14 specifically regulated MDA5-dependent IFN-β production by overexpressing MDA5 or RIG-I in A549 cells, which is known to induce IFN expression. IFN-β production due to exogenous expression of RIG-I remained unaffected following PARP14i treatment. However, there was a significant ∼3-fold reduction in IFN-β production following PARP14i treatment in MDA5 expressing cells **(Fig. 3B)**. To corroborate these findings we use two different viruses, Sendai virus (SeV) and encephalomyocarditis virus (EMCV), that are specific agonists of RIG-I and MDA5, respectively. PARP14i treatment reduced IFN-β production ∼3-fold during EMCV infection but had no effect on SeV induced IFN production in BMDMs **(Fig. 3C)** and A549 cells **(Fig. 3D)**. These results indicate that PARP14 promotes MDA5 but not RIG-I dependent IFN-β production.

**Fig. 3.**
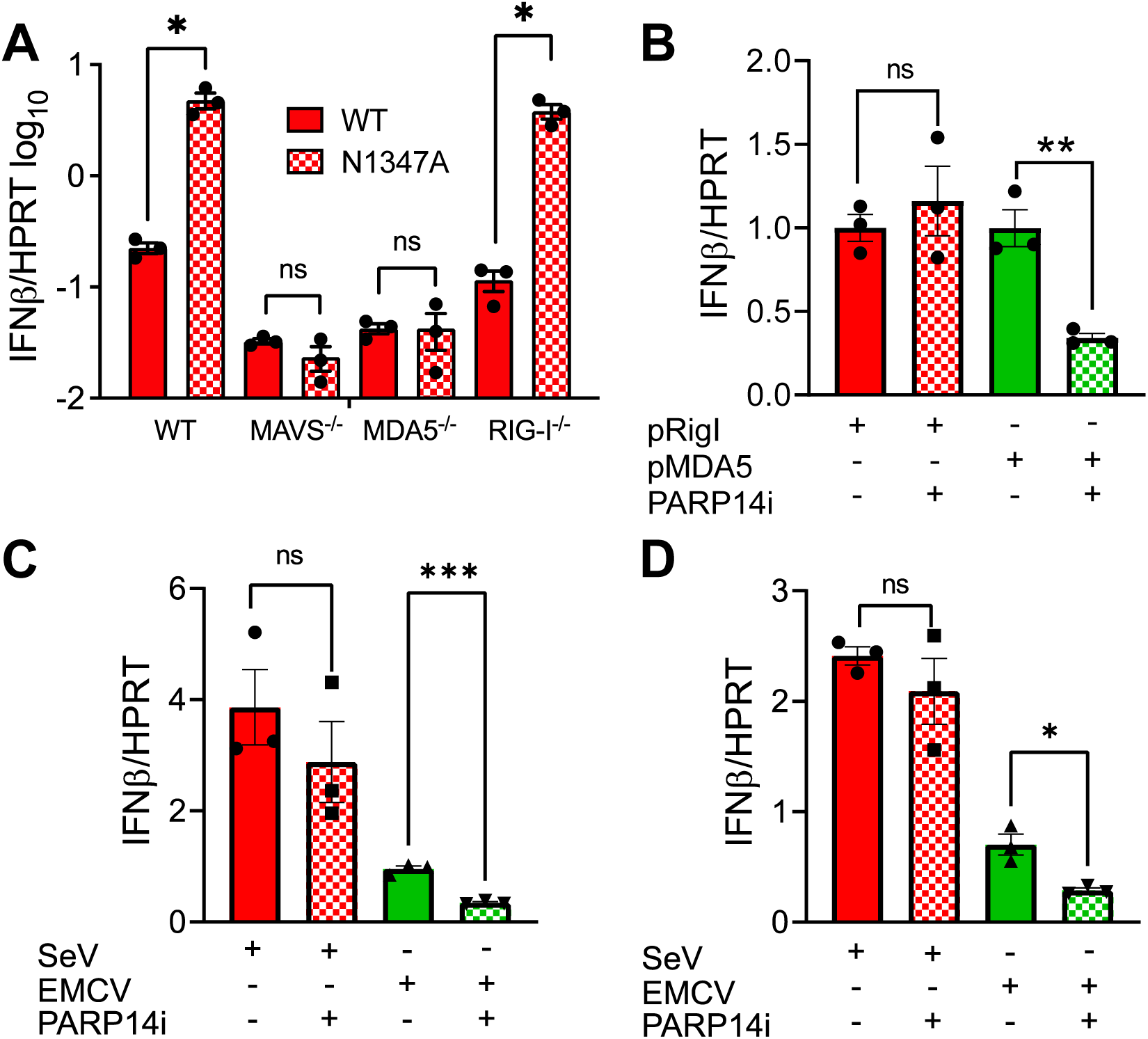
PARP14 promotes MDA-5 dependent IFN-*β* production. A) Bone-marrow derived cells were harvested from WT, MAVS^-/-^, MDA-5^-/-^, and RIG-I^-/-^ mice and differentiated into M0 macrophages. Then they were infected with MHV WT or N1347A at an MOI of 0.1. RNA was isolated from cells at 12 hpi and IFNβ mRNA was quantified by qPCR using ΔCt method. The results in A are from one experiment representative of two independent experiments each with N=3 biological replicates per group in each experiment. B) A549 cells were transfected with 0.5 *μ*g/ml of indicated plasmid. Then either DMSO or 100 nM PARP14i were added to the cells 4 hours post-transfection (hpt). RNA was isolated from cells at 24 hpt and IFNβ mRNA was quantified by qPCR using ΔCt method. C-D) BMDMs (C) or A549 cells (D) were infected with either SeV at an MOI of 1 or EMCV at an MOI of 2. 100 nM PARP14i or DMSO was then added 1 hpi. RNA was isolated from cells at 24 hpi (A549 cells) or 36 hpi (BMDMs) and IFNβ mRNA was quantified by qPCR using ΔCt method. The results in B-D are from one experiment representative of three independent experiments each with N=3 biological replicates per group for each experiment.

### RNAseq analysis identifies over 100 genes up-regulated by PARP14

Next, we used RNAseq to identify all genes that are influenced by PARP14-dependent MARylation, in addition to IFNs and ISGs. To identify PARP14 regulated genes, we analyzed the transcriptome of BMDMs that are infected by N1347A with and without PARP14i. As expected from previous studies, N1347A infected DMSO-treated BMDMs had elevated levels of IFN-I, ISGs, and dozens of other genes involved in antiviral immunity. Upon treatment with PARP14i, the mRNA levels of a substantial number of these genes were reduced, indicating that these mRNAs are increased by PARP14-dependent MARylation. Overall, we found that 622 genes were upregulated by PARP14, while 355 genes were downregulated, for a total number of 977 differentially expressed genes expressed above a threshold fold change (FC) of 1.5-fold and threshold P-adjusted value of 0.001 **(Fig. 4A)**. Next, we performed a functional analysis on the genes that are regulated by PARP14 MARylation activity. Many of the genes promoted by PARP14 were genes involved in antiviral immunity **(Fig 4B)**. However, genes involved in apoptosis, transcription, and ubiquitin ligase conjugation were also upregulated. Representative genes from each of these pathways were validated by qPCR **(Fig. 4C)**. Furthermore, we took a closer look at some of the genes observed to be upregulated or downregulated over 20-fold. These genes were found to cluster in biological processes of transcription and transcriptional regulation with very low p-value (upon functional enrichment analysis by DAVID). Overall, this data suggests that PARP14 is an important regulator of antiviral immunity but may also regulate other cellular processes such as cell death and transcription.

**Fig. 4.**
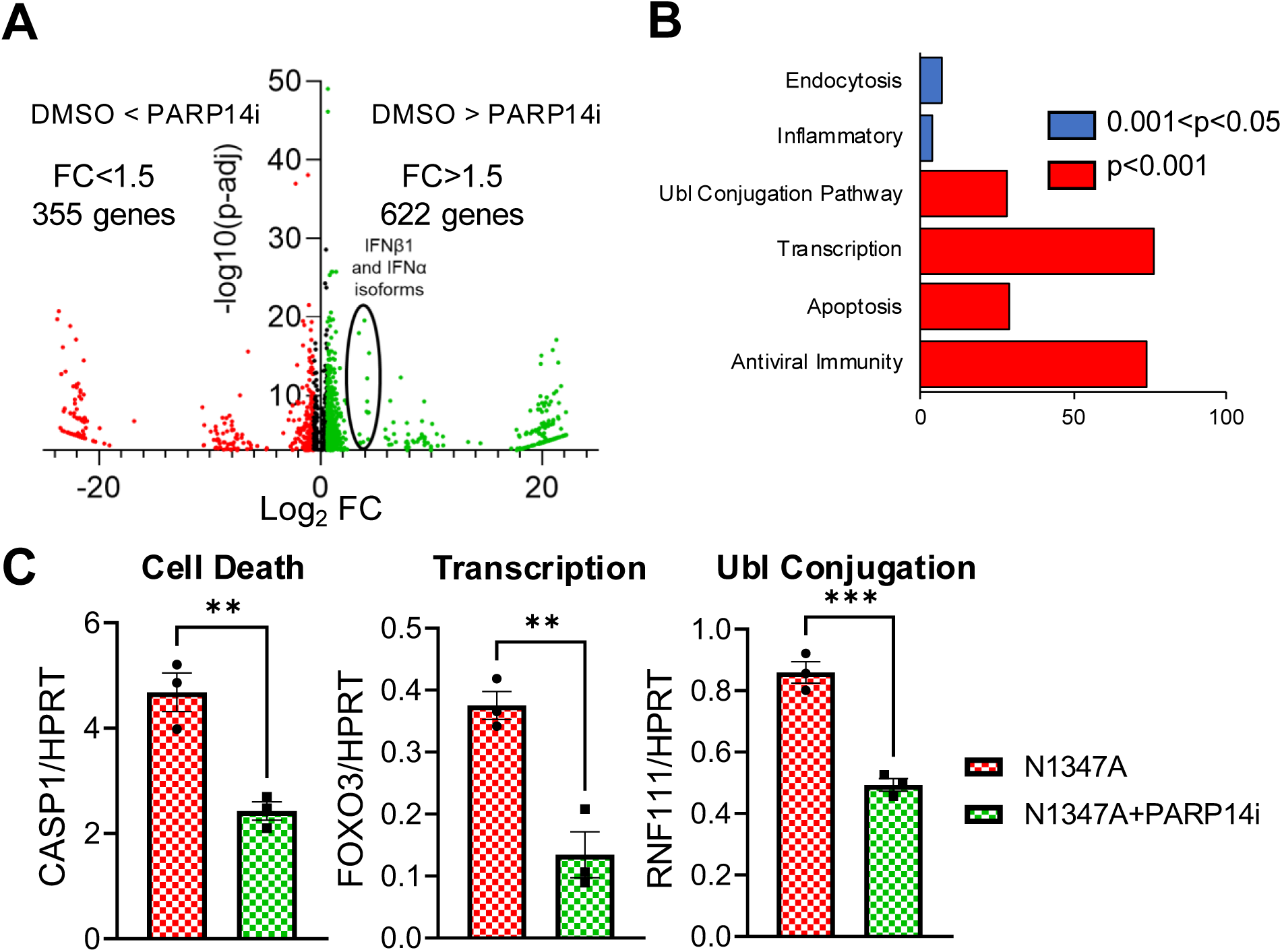
PARP14 regulates the transcription of dozens of genes with multiple biological functions following MHV-N1347A infection of BMDMs. A) WT BMDMs were infected with N1347A MHV at an MOI of 0.1 and then treated with DMSO or 100 nM PARP14i at 1 hpi. RNA was collected from these cells at 12 hpi and RNAseq was performed to quantify the transcriptome with and without PARP14i. The ratio of gene expression in the presence or absence of PARP14i treatment was plotted against minus log_10_ value of the p-adjusted value as a volcano plot. B) Gene ontology analysis/pathway enrichment analysis of genes that were identified to be promoted by PARP14 catalytic activity carried out using DAVID gene ontology tool. Pathways enriched with a p-value of less that 0.05 were plotted. C) WT BMDMs were infected with MHV N1347A at an MOI of 0.1 and treated with DMSO or 100 nM PARP14i at 1 hpi. RNA was isolated from cells at 12 hpi and CASP1, FOXO3 and RNF111 mRNAs were quantified by qPCR using ΔCt method. RNAseq data shown in A-B are from 1 experiment with N=3. Data shown in C are from 1 experiment and are representative of 2 independent experiments with N=3 biological replicates per group for each experiment.

### PARP14 is required for the repression of Mac1-mutant coronavirus replication

We previously found that *parp14* knockdown mildly increased the replication of MHV-N1347A in BMDMs, however congenital *parp14* KO mice did not have increased MHV-N1347A replication [3]. To determine if PARP14 was required to repress MHV-N1347A, we infected our *parp14^-/-^* and *parp14^+/+^* BMDMs as described above. As previously observed, MHV-N1347A had a significant growth defect compared to MHV-WT virus in *parp14+/+* cells, regardless of whether the cells were or were not treated with tamoxifen **(Fig. 5A)**. Interestingly, *parp14^-/-^* mostly restored the replication of MHV-N1347A, suggesting that PARP14 is required for restricting MHV-N1347A virus replication in BMDMs **(Fig. 5B)**. We also found that MHV-N1347A replication was increased to levels similar to those of MHV-WT in *parp14^-/-^* bone-marrow-derived dendritic cells (BMDCs) **(Fig. 5C)**. To test whether this increased MHV-N1347A replication in *parp14^-/-^*cells is due to PARP14-dependent MARylation, we measured MHV-N1347A replication in the presence and absence of PARP14i. In the presence of PARP14i, N1347A replication was similarly restored to near WT levels, suggesting that the catalytic activity of PARP14 was required for restricting MHV-N1347A replication **(Fig. 5D)**.

**Fig. 5.**
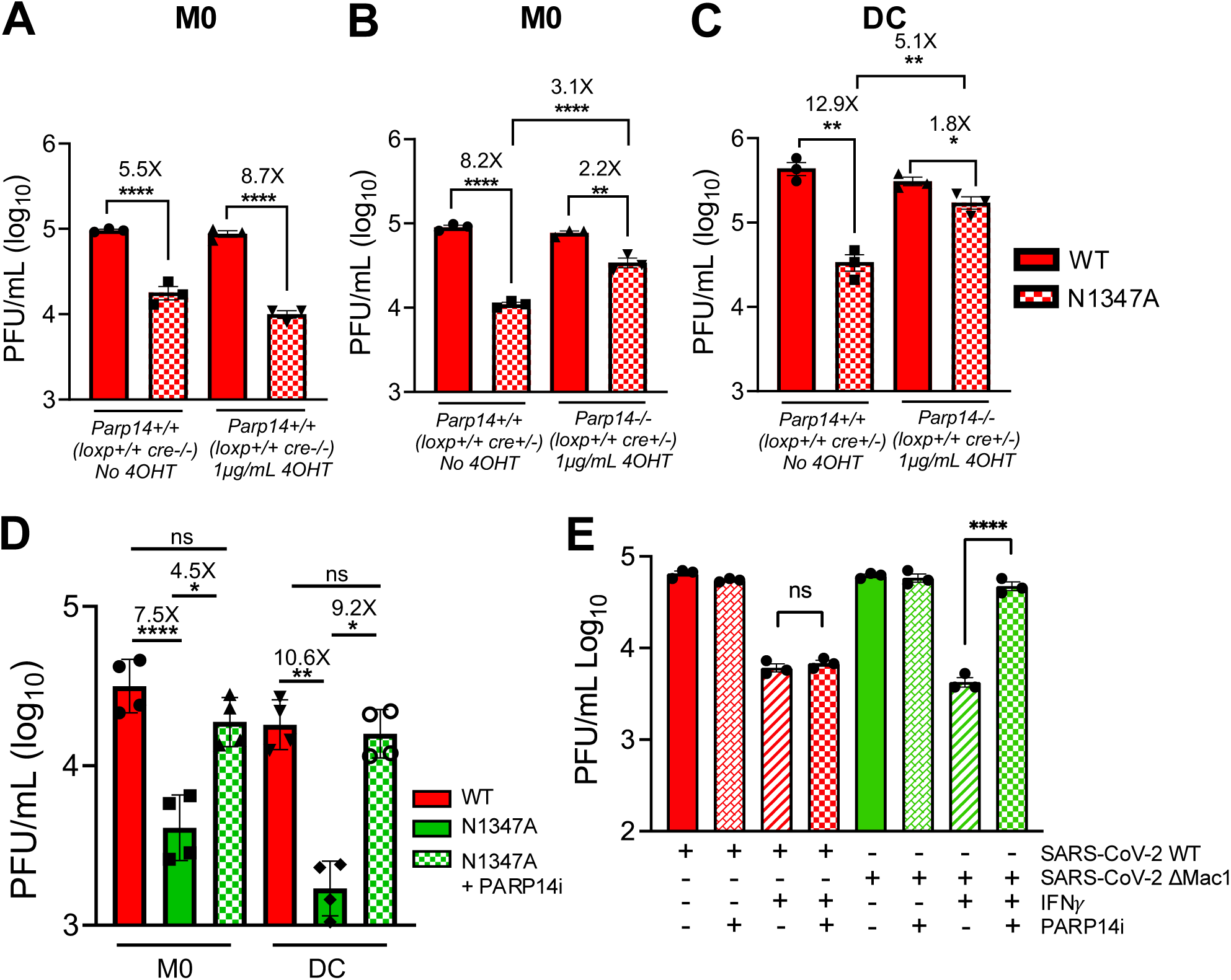
PARP14 inhibits the replication of Mac1-mutant MHV and SARS-CoV-2. A) *Parp14* floxed *cre+/-* BMDMs were treated with and without 4-OHT. 24 hours later these cells were infected with WT and N1347A MHV at an MOI of 0.1. At 20 hpi cells and supernatants were collected and progeny virus was quantified by plaque assay. B-C) PARP14 floxed *cre+* BMDMs (B) and DCs (C) were treated with or without 4-OHT. 24 hours later these cells were infected with WT and N1347A MHV at an MOI of 0.1. At 20 hpi cells and supernatants were collected and progeny virus was quantified by plaque assay. Data shown in A-C are from 1 experiment and are representative of 3 independent experiments with N=3 biological replicates per group for each experiment. D) PARP14^+/+^ BMDMs and DCs were infected with WT and N1347A MHV at an MOI of 0.1 and then treated with DMSO or PARP14i (1 *μ*M). At 20hpi cells and supernatants were collected and progeny virus was quantified by plaque assay. Data shown in D are from 1 experiment and are representative of 3 independent experiments with N=4 biological replicates per group for each experiment. E) WT Calu-3 cells were either mock-treated or IFN*γ*-treated (50 Units) O/N. These cells were then infected with WT and *Δ*Mac1 SARS-COV-2 at an MOI of 0.1 and treated with DMSO or PARP14i (1 *μ*M) at 1 hpi. At 48 hpi cells and supernatants were collected and progeny virus was quantified by plaque assay. Data shown in E are from 1 experiment and are representative of 3 independent experiments with N=3 biological replicates per group for each experiment.

We next tested whether PARP14 was required to restrict the replication of severe acute respiratory syndrome coronavirus 2 (SARS-CoV-2) ΔMac1. We previously found that SARS-CoV-2 ΔMac1 replicates near SARS-CoV-2 levels in Calu-3 cells, but that SARS-CoV-2ΔMac1 is more sensitive to IFN-γ pre-treatment than WT virus, and IFN-γ induces the expression of several PARPs, including PARP14 [25]. To determine if PARP14 is required for IFN-γ mediated restriction of SARS-CoV-2ΔMac1, we infected IFN-γ pretreated Calu3 cells with SARS-CoV-2 WT and ΔMac1 with and without PARP14i. Upon treatment with PARP14i, we observed that WT virus was not impacted either in the presence or absence of IFN-γ, in contrast, the replication of SARS-CoV-2ΔMac1 in cells pre-treated with IFN-γ was significantly increased to levels near that of non-IFN-γ treated cells **(Fig. 5E)**. Overall, these results suggest that MARylation triggered by PARP14 is required to restrict the replication of Mac1-deficient MHV and SARS-CoV-2 viruses.

### PARP14 restricts HSV-1 replication

PARP14 is one of several PARPs that has evolved under positive selection in primates, indicating that it is involved in host-pathogen conflict [22]. Thus, we hypothesized that in addition to CoVs, it might repress the replication of other viral families. First, we tested the effect of PARP14 deletion on HSV-1, which has been co-evolving with humans for several million years. Furthermore, PARP14 was identified as a host factor that binds to HSV-1 DNA [26, 27]. We infected A549 WT and two different clones of PARP14 KO cells [3] with HSV-1 (strain KOS) at a low MOI (0.01PFU/mL) and found that replication of HSV-1 was enhanced by over 2-logs throughout the infection in PARP14 KO cells, strongly indicating that PARP14 represses HSV-1 replication **(Fig. 6A)**. HSV-1 infection is characterized by activation of some of its immediate early genes like ICP0, ICP4 and ICP27. As a separate measure of replication, we infected WT and KO cells with 0.1 MOI HSV-1 and quantified the expression of ICP0, ICP4 and ICP27 mRNA transcripts at 24 hpi. We observed increased gene expression in PARP14 KO cells compared to WT cells, again indicating that PARP14 restricts HSV-1 infection **(Fig. 6B).** We then corroborated this observation by quantifying ICP0, ICP4 and ICP27 mRNA levels in A549 cells infected with HSV-1 and treated with PARP14 degron (DEG) or its corresponding negative control (CTL). Again, we saw an increase in the transcripts in DEG treated cells, confirming that the observed increase in expression of ICP genes was due to the absence of PARP14 **(Fig. 6C)**. Next, we tested the ability of HSV-1 to form plaques on A549 WT and PARP14 KO cells using a plaquing efficiency assay. Similar to previous observations, we found that PARP14 KO also enhanced the plaquing efficiency of HSV-1 compared to WT cells by >2-logs on average **(Fig. 6D)**. Since the MARylating activity of PARP14 was responsible for restricting CoV replication we tested the ability of PARP14i to enhance HSV-1 infection. Interestingly, HSV-1 replication was not affected in PARP14i treated A549 cells compared to untreated cells, suggesting that PARP14 restricted HSV-1 replication in a MARylation-independent manner **(Fig. 6E)**. Overall, these data show that PARP14 restricts HSV-1 infection and does so in an ADP-ribosylation independent manner.

**Fig. 6.**
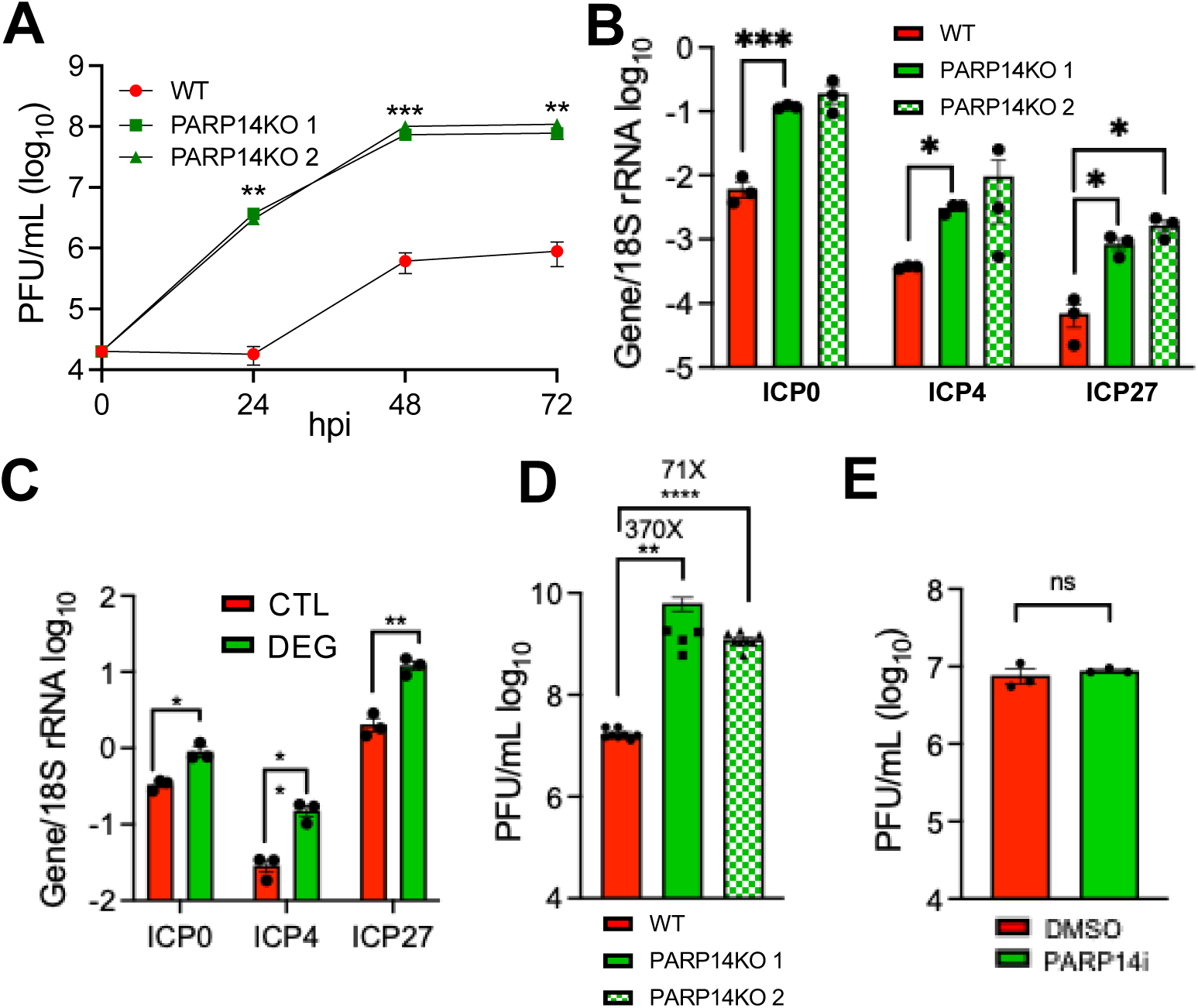
PARP14 restricts the replication of HSV-1 in A549 cells. A) WT and PARP14 KO A549 cells were infected with HSV-1 at an MOI of 0.01 PFU/cell. Cells and supernatants were collected at indicated time points and progeny virus was quantified by plaque assay. B) WT and PARP14 KO A549 cells were infected with HSV-1 at an MOI of 0.1. At 24 hpi RNA was harvested from infected cells and HSV-1 immediate early gene ICP4, ICP0 and ICP27 mRNAs were quantified by qPCR using ΔCt method having 18S rRNA abundance as the endogenous control. The data in A-B is from 1 experiment representative of 2 independent experiments. N=3 biological replicates per group for each experiment. C) WT A549 cells were infected with HSV-1 at an MOI of 0.01 PFU/cell. At 1 hpi cells were treated with 1 μM PARP14 Degron control (CTL) or PARP14 Degron (DEG). At 24 hpi RNA was harvested from infected cells and HSV-1 immediate early gene ICP4, ICP0 and ICP27 mRNAs were quantified by qPCR using ΔCt method having 18S rRNA abundance as the endogenous control. The data is from 1 experiment representative of 2 independent experiments. N=3 biological replicates per group. D) WT and PARP14 KO A549 cells were infected with serial dilutions of HSV-1 starting at a concentration of 1.85×10^9^ PFU/ml and plaquing efficiency was determined at 24 hpi. The data in E the combined results of 3 separate experiments, N=9 biological replicates per group. E) WT A549 cells were infected with HSV-1 at an MOI of 0.1 PFU/cell and then treated with DMSO or PARP14i (1 *μ*M) at 1 hpi. Cells and supernatants were collected at 24 hpi and progeny virus was quantified by plaque assay. The data in E is from 1 experiment representative of 2 independent experiments. N=3 biological replicates per group for each experiment.

### PARP14 promotes the replication of vesicular stomatitis virus (VSV)

Next, we tested the ability of PARP14 to restrict the replication of additional types of RNA viruses. First, we tested the infectivity of lymphocytic choriomeningitis virus clone-13 (LCMV-cl13), an ambisense RNA virus, in WT and PARP14 KO BMDMs. To measure infectivity of LCMV-cl13 in BMDMs, we measured the amount of LCMV nucleoprotein (LCMV-NP) using a fluorescently labeled antibody against LCMV-NP (clone anti-VL4) and intracellular flow cytometry. PARP14 KO cells had comparable levels of LCMV-NP+ cells compared to WT cell, ∼4-5%. These data indicate that LCMV-cl13 infection is not impacted by PARP14 **(Fig. S3A-B)**. To further explore the role of PARP14 in the replication of negative-sense RNA viruses, we tested the ability of VSV-GFP to replicate in PARP14 KO A549 cells. Interestingly, VSV-GFP replication was significantly reduced by in A549 PARP14 KO cells compared to WT cells, indicating that PARP14 promotes VSV-GFP replication **(Fig. 7A)**. To confirm this result, we created a new set of A549 WT and PARP14 KO cells using separate control and PARP14 targeting guide RNAs **(Fig. S4)**. Again, we found that VSV replication was significantly reduced in these cells **(Fig. 7B)**. Furthermore, the addition of the PARP14 degrader (DEG) significantly reduced the replication of VSV in WT cells, while the CTL compound had no effect **(Fig. 7B)**. To determine if these differences could be due to an increase in IFN-levels, we measured the level of IFN following infection with VSV in both PARP14 KO and PARP14i treated cells. We consistently found no substantial differences in IFN production following VSV infection in PARP14 depleted cells **(Fig. S5A)**. To further confirm that there was no impact of IFN on virus replication, we treated A549 cells with a JAK inhibitor (JAKi) that has previously been used to ablate the production of ISGs [28, 29]. While the JAKi significantly reduced the levels of PARP14, an ISG **(Fig. S5B)**, it did not lead to increased replication of VSV in WT or PARP14 KO cells **(Fig. S5C)**. Finally, we tested whether the ADP-ribosyltransferase activity of PARP14 supported VSV replication by treating A549 WT cells with PARP14i. PARP14i treatment did not significantly reduce VSV replication, indicating that the proviral function of PARP14 was largely independent of its MARylating activity **(Fig. 7C)**. Overall, these results suggest that PARP14 can inhibit the replication of CoVs and HSV-1 but enhances the replication of VSV, a negative-sense RNA virus.

**Fig. 7.**
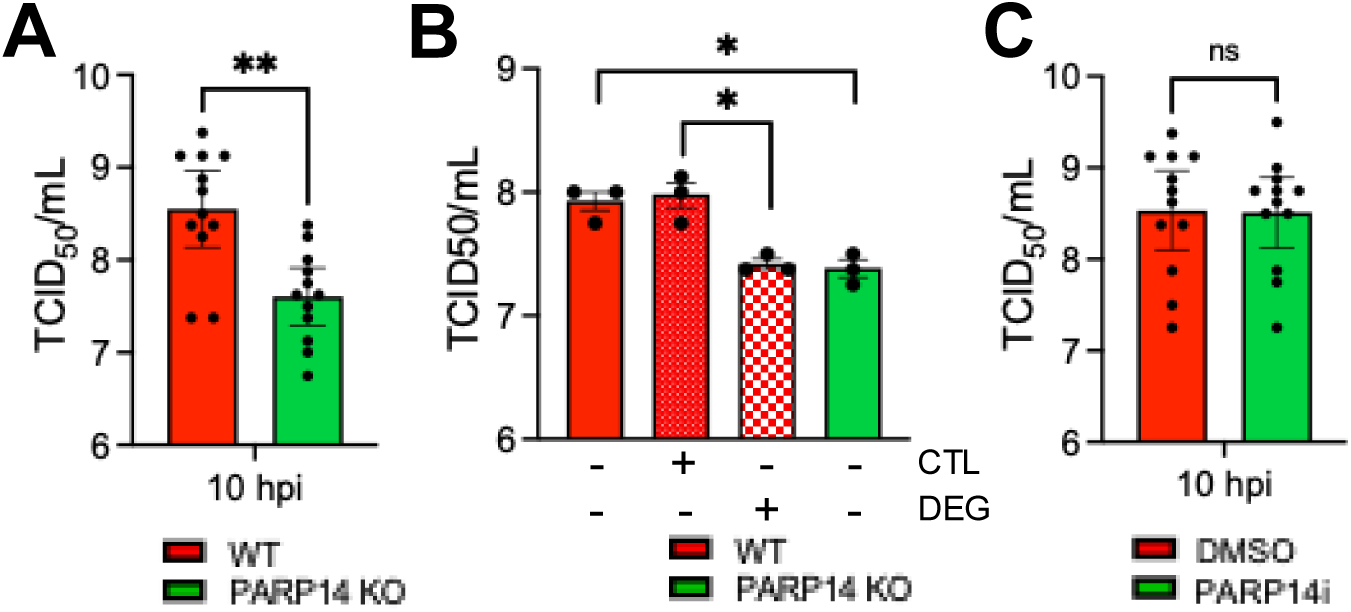
PARP14 is required for efficient replication of VSV-GFP in cell culture. A) A549 WT and PARP14 KO cells were infected with VSV-GFP at 1 MOI. At 10 hpi, cells and supernatants were collected and progeny virus was quantified by TCID_50_. Results in A are the combined results of 4 independent experiments. N=12 biological replicates. B) A549-ACE2 WT and PARP14 KO cells were infected with VSV-GFP at 1 MOI and then treated with 1 μM PARP14 Degron control (CTL) or PARP14 Degron (DEG) at 1 hpi. At 10 hpi, cells and supernatants were collected and progeny virus was quantified by TCID_50_. Results in B are from one experiment representative of 2 independent experiments. N=3 biological replicates per group for each experiment. C) A549 WT cells were infected with VSV at an MOI of 1 and then treated with DMSO or 100 nM PARP14i (P14i) at 1 hpi. At 10 hpi, cells and supernatants were collected and progeny virus was quantified by TCID_50_. Results in C are the combined results of 4 independent experiments. N=12 biological replicates.

## DISCUSSION

PARP14 has emerged as a critical host factor that impacts multiple immune signaling pathways. It was initially found to bind STAT6 and promote IL4-dependent anti-inflammatory gene regulation [15]. This function of PARP14 promotes T_H_2 gene expression and IgE production in immune cells [30]. Recently, a PARP14 inhibitor (RBN-3143) was reported to mitigate the effects of allergen-induced IgE production and overproduction of mucus, reduced T_H_2 cell abundance and restored *α*PD-1 sensitivity to tumors [31]. PARP14 also inhibits STAT-1 phosphorylation by MARylating key residues, and knockdown or knockout of PARP14 increased pro-inflammatory cytokines in primary human and mouse M1, but not M2 macrophages [16]. In contrast, PARP14 promoted IFN-I and pro-inflammatory cytokine production in RAW 264.7 cells following LPS treatment and *S. typhimurium* infection [17]. PARP14 also promoted IFN-I production in response to MHV infection and poly(I:C) stimulation in BMDMs and A549 cells, respectively [3]. In this study, we demonstrated that the enzymatic activity of PARP14 promotes MDA5-dependent IFN-I and IFN-III production when stimulated with a dsRNA analog poly(I:C), EMCV, or CoV infection by directly quantifying IFN-I and IFN-III mRNA and protein levels in multiple cell lines. In general, the impact of PARP14 on innate and adaptive immune pathways is cell type and condition specific. It remains unclear if PARP14 both enhances IFN-I and IFN-III production and limits IFN signaling in the same cells, or if it only represses IFN signaling in M1 macrophages. It is feasible that PARP14 may dampen IFN signaling to control its own ability to stimulate IFN production. How PARP14 regulates MDA-5-dependent IFN production and the broader impacts of these functions remains unclear and is an active topic of research.

In addition to inducing IFN, PARP14 expression is stimulated by IFNs and virus infections and that its ADP-ribosyltransferase activity triggers the production of IFN-induced ADP-ribose bodies (ICABs) [3]. ICABs are regions of the cytoplasm with punctate ADP-ribose signal that has been observed by immunofluorescence following IFN-*γ* or poly(I:C) treatment [18-21, 32]. In addition, increased mono-ADP-ribosylation is visible by Western blotting following the same treatments in A549 cells [18]. Importantly both signals were dependent on the catalytic activity of PARP14 and can be reversed by macrodomain proteins, including the SARS-CoV-2 Mac1 protein [18, 32]. These bodies also contain PARP9, DTX3L, and p62, and also overlap with ubiquitin staining. Furthermore, DTX3L and p62 are ADP-ribosylated by PARP14, likely within ICABs, and are required for the maintenance of the ICABs [18, 20, 21, 33]. Despite this, there is no current data showing that PARP14 restricts virus infection. Here, we found that the catalytic activity of PARP14 was required to repress the replication of Mac1-mutant CoVs. Furthermore, our findings that PARP14 catalytic activity is required for the repression of SARS-CoV ΔMac1 only in the presence of IFN-*γ* strongly suggests that ICABs play a key role in the repression of CoV replication. Future studies will determine if p62, DTX3L, or other PARP14 targets within ICABs participate in the restriction of CoV replication and the mechanisms by which they do so.

PARP14 is one of 4 PARPs that has evolved under positive selection, further indicating that it is likely involved in host-pathogen conflict [22]. In addition to restricting Mac1-mutant CoVs, we also found that it was required for repression of HSV-1. HSV-1 and other herpesviruses have been evolving with mammals for millions of years, so it is maybe more likely that PARP14 is co-evolving with herpesviruses than CoVs. Finally, we found that PARP14 enhances the replication of VSV in an ADP-ribosylation independent manner. PARP14 has been shown to interact with SARS-CoV-2 RNA [35] and HSV-1 DNA [26], thus the mechanism by which PARP14 restricts and promotes virus replication could be through interactions with nucleic acids. In fact, PARP14 bound to the HSV-1 genome, in association with other DNA repair proteins, as early as 3 hours post-infection, suggesting that PARP14 possibly binds to DNA to inhibit transcription and/or DNA replication [26, 27]. These interactions may help explain the positive selection of PARP14 as it may need to rapidly evolve to mutations in DNA or RNA sequences within viral genomes.

PARP14 is a large protein (203kDa) with multiple domains [12]. The ADP-ribosyl transferase (ART) domain of PARP14 binds to NAD^+^ and transfers an ADP-ribose moiety to its targets. Its ART domain has an HYL (hydrophobic residue-tyrosine-leucine) domain as part of the catalytic triad, which restricts its ART activity to MARylation [36, 37]. PARP14 also has 3 tandem macrodomains (MD), of which MD1 has ADP-ribosylhydrolase (ARH) activity against MARylated protein and nucleic acid substrates, making it a rare enzyme that both adds and reverses the same post-translational modification [13, 14]. PARP14 also has a WWE domain that can bind to ADP-ribose subunits [38], several KH domains that facilitate protein and nucleic acid binding, and several RNA recognition motifs (RRMs) [12]. However, the roles and substrates of these domains, especially during virus infection, remain largely unknown. Given the potential of PARP14 to bind and modify protein and nucleic acid substrates through its many domains, identifying the protein and/or nucleic acid binding partners of PARP14 during infection and the domains required for these interactions could reveal several novel mechanisms by which PARP14 regulates virus replication and pathogenesis.

## MATERIALS AND METHODS

### Biosafety statements

All work with SARS-CoV-2 was conducted in the University of Kansas EHS-approved BSL-3 facility following standard operating procedures. All work with wild-type and recombinant Ebola virus and Nipah virus was performed in the biosafety level 4 (BSL-4) facility of Boston University’s National Emerging Infectious Diseases Laboratories (NEIDL) following approved standard operating procedures in compliance with local and national regulations pertaining to handling BSL-4 pathogens and Select Agents. Animal studies were approved by the Institutional Animal Care and Use Committees at the University of Iowa and the University of Kansas and met stipulations of the Guide for the Care and Use of Laboratory Animals.

### Cell culture and reagents

NHDF, 293T, Hela-MVR, A549, A549-PARP14KO, A549-ACE2 (a generous gift from Susan Weiss, University of Pennsylvania), BHK, 17Cl-1, and Vero E6 cells were grown in Dulbecco’s modified Eagle medium (DMEM) supplemented with 10% fetal bovine serum (FBS). Calu-3 cells (ATCC) were grown in MEM supplemented with 20% FBS. Baby hamster kidney cell line BSR-T7/5 constitutively expressing T7 RNA polymerase was maintained in Glasgow’s Minimum Essential Medium (G-MEM) with 10% FBS. Bone marrow derived cells were grown in RPMI supplemented with 10% FBS and differentiated into M0 macrophages using M-CSF (Millipore-Sigma) and dendritic cells (DCs) using GM-CSF (Millipore-Sigma). All cell lines were grown at 37℃ and 5% CO_2_. IFN-*γ* was purchased from R&D Systems and Poly(I:C) HMW (0.5μg/mL) was purchased from Millipore-Sigma. RBN012759 (PARP14i) was purchased from Atomwise. RBN012811 (DEG) and its associated negative control RBN013527 (CTL) were generously provided by Dr. Mario Niepel (Ribon Therapeutics). Cells were transfected with either Polyjet (Amgen), Lipofectamine 3000 (Fisher Scientific), or jetOPTIMUS (Polyplus) per the manufacturer’s instructions.

### Creation of inducible PARP14 knock-out mice

*Animals.* The insertion of loxP sites flanking exon 2 of PARP14 was carried out by the University of Iowa genome editing facility. C57BL/6J mice were purchased from Jackson Labs (000664; Bar Harbor, ME). Male mice older than 8 weeks were used to breed with 3-5-week-old super-ovulated females to produce zygotes for pronuclear injection. Female ICR (Envigo; Hsc:ICR(CD-1)) mice were used as recipients for embryo transfer. All animals were maintained in a climate-controlled environment at 25℃ and a 12/12 light/dark cycle.

*Preparation of Cas9 RNPs and the injection mix.* Chemically modified CRISPR-Cas9 crRNAs targeting regions 5’ and 3’ of exon 2 of PARP14 **(Table S1)** and CRISPR-Cas9 tracrRNA were purchased from IDT (Alt-R® CRISPR-Cas9 crRNA; Alt-R® CRISPR-Cas9 tracrRNA (Cat# 1072532)). The crRNAs and tracrRNA were suspended in T10E0.1 and combined to 1 ug/ul (∼29.5 uM) final concentration in a 1:2 (μg:μg) ratio. The RNAs were heated at 98℃ for 2 min and allowed to cool slowly to 20℃ in a thermal cycler. The annealed cr:tracrRNAs were aliquoted to single-use tubes and stored at -80℃. Cas9 nuclease was also purchased from IDT (Alt-R® S.p. HiFi Cas9 Nuclease). Cr:tracr:Cas9 ribonucleoprotein complexes were made by combining Cas9 protein and cr:tracrRNA in T10E0.1 (final concentrations: 250 ng/*μ*l (∼1.6 *μ*M) Cas9 protein and 200 ng/*μ*l (∼6.1 *μ*M) cr:tracrRNA). The Cas9 protein and annealed RNAs were incubated at 37℃ for 10 minutes. The RNP complexes were combined with single-stranded repair template, containing the loxP sites to be inserted around exon2, and incubated an additional 5 minutes at 37℃ (single-stranded repair template sequence available upon request). The concentrations in the injection mix were 50 ng/*μ*l (∼0.3 *μ*M) Cas9 protein and 20 ng/*μ*l (∼0.6 *μ*M) each cr:tracrRNA and 40 ng/*μ*l single-stranded repair template.

*Collection of embryos and injection.* Pronuclear-stage embryos were collected using methods described in [39]. Embryos were collected in KSOM media (Millipore; MR101D) and washed 3 times to remove cumulous cells. Cas9 RNPs and double-stranded repair template were injected into the pronuclei of the collected zygotes and incubated in KSOM with amino acids at 37℃ under 5% CO_2_ until all zygotes were injected. The CRISPR injection protocol was previously described[40]. Fifteen to 25 embryos were immediately implanted into the oviducts of pseudo-pregnant ICR females. Offspring were heterozygote for the loxP insertions. These mice were sequentially bred to result in floxed homozygote mice. Genotyping was performed using PCR amplification with primers listed in **Table S2**. and restriction digestion by EcoR1. These were then crossed with C57BL/6.Cg-Ndor1^Tg(UBC-cre/ERT2)1Ejb^/IJ mice (Jax strain # 007001) till mice colonies that were homozygous for *loxP* sites and heterozygous for *cre* gene were confirmed by genotyping. The resulting mice were backcrossed twice before using these mice for the experiments described in this study.

### Creation of PARP14 knock-out A549-ACE2 cells

A549-ACE2 cells were cultured in DMEM supplemented with 10% FBS at 37°C with 5% CO₂. For CRISPR/Cas9 transfection, cells were seeded in 6-well plates at 2 × 10⁵ cells per well, before transfection. At 40-80% confluency, cells were transfected with either PARP14 CRISPR/Cas9 KO plasmid (sc-402812, Santa Cruz Biotechnology) or control CRISPR/Cas9 plasmid (sc-418922, Santa Cruz Biotechnology) using PolyJet transfection reagent according to manufacturer’s instructions. After 72 hours of incubation, successful transfection was confirmed by GFP expression via fluorescence microscopy. GFP-positive cells were isolated using a BD FACSymphony cell sorter, and single cells were sorted into individual wells to establish monoclonal populations. WT control and PARP14 knockout expanded clones were validated by Western blot analysis using anti-PARP14 mouse monoclonal antibody (sc-377150, Santa Cruz Biotechnology).

### Virus infections

*MHV-JHM^IA^*. Recombinant WT and N1347A MHV JHM^IA^ virus used in this study have been previously reported [3, 41, 42]. Virus stocks were grown in 17Cl-1 cells and titered by plaque assay in Hela-MVR cells. BMDMs and DCs were infected at an MOI of 0.1 PFU/cell for indicated periods of times after a 60-min adsorption phase. Virus titers were determined by plaque assay on Hela-MVR cells.

*SARS-CoV-2*. Recombinant SARS-CoV-2 WT and ΔMac1 virus based on the original Wuhan isolate used in this study has been previously reported [25]. Virus stocks were grown and titered by plaque assay in VeroE6 cells. A549-ACE2 cells were infected at an MOI of 0.1 for 48 hours after a 60-min adsorption phase. Calu3 cells were infected at an MOI of 0.1 for 48 hours after a 120-min adsorption phase and Trypsin-TCPK treatment (1μg/mL). Virus titers were determined in Vero E6 cells using plaque assay. All the SARS-CoV-2 experiments were carried out under BSL-3 conditions at the University of Kansas, following approved SOPs.

*HSV-1.* Stocks of HSV-1 strain KOS was propagated and titered as previously described [43]. For viral yield assays, A549 and A549-PARP14KO cells (5×10^4^) were seeded in triplicate in a 24-well plate and infected at an MOI of 0.1 PFU/cell with HSV-1. At 1 hpi, cells were washed twice with PBS to remove unabsorbed virus. Infected cells were harvested and collected 24 hpi and titered by standard plaque assays on Vero cells. For PARP14 inhibitor experiments, A549 cells were treated with or without PARP14 inhibitor, and infections were carried out as described for the HSV-1 yield assays. To determine plaquing efficiencies, A549 and A549-PARP14KO cells were plated in triplicate in 12-well plates and infected with serial dilutions from HSV-1 stocks, and titers were determined by standard plaque assays. To assess virus transcript levels, A549 cell lines (5×10^4^) were plated in triplicate in a 24-well plate. Cells were infected at an MOI of 0.1 with HSV-1 and washed twice with PBS 1 hpi. At 6 hpi, media were removed, and cells were resuspended in TRIzol reagent (250 μL). RNA was isolated according to manufacturer’s protocol and reverse transcribed as published (ref). cDNAs of viral and cellular transcripts were amplified using primers for ICP0, ICP4, ICP27, and 18S rRNA as previously described [44].

*VSV*. VSV-GFP Indiana strain was generously provided by Dr. Asit Pattnaik (University of Nebraska-Lincoln) [45]. Virus stocks were grown and titered by plaque assay in VeroE6 cells. A549 cells were infected with VSV-GFP at an MOI of 1 PFU/cell. Infected cells were then incubated and collected at 10 hpi after a 60-min adsorption phase. Virus titer was determined by 50% tissue culture infectious dose (TCID_50_) assay in Vero E6 cells.

*LCMV*. LCMV Clone-13 (LCMV-cl13) virus stocks were grown in BHK cells and titered by plaque assay in Vero E6 cells. BMDM cells were infected with LCMV-cl13 at an MOI of 2-6. Infected cells were then incubated and collected at 24 hpi. For surface staining, samples were treated with Fc block (CD16/32, eBioscience) and then incubated with Tonbo Ghost Viability dye (violet 510, Tonbo biosciences), F4/80 (Pacific orange, Invitrogen, CLONE) and LCMV-NP (clone VL4 (BioXcell), self-conjugated to Alexa Fluor 488 following manufacturer’s instructions, (Thermo Fisher Scientific)). All flow cytometry data were analyzed using FlowJo software (BD Biosciences).

### Immunoblotting

Total cell extracts were lysed in sample buffer containing SDS, protease and phosphatase inhibitors (Roche), *β*-mercaptoethanol, and a universal nuclease (Fisher Scientific) and boiled at 95°C for 5 minutes. Proteins were resolved on an SDS polyacrylamide gel, transferred to a polyvinylidene difluoride (PVDF) membrane, hybridized with a primary antibody, reacted with an infrared (IR) dye-conjugated secondary antibody, visualized using a Li-COR Odyssey Imager (Li-COR), and analyzed using Image Studio software. Primary antibodies used for immunoblotting included anti-SARS-CoV-2 N (SinoBiological 40143-R001, 1:5000), anti-OAS3 (Cell Signaling 41440, 1:250), anti-PARP14 (Santa Cruz Biotechnology SC-377150, 1:100), and anti-GAPDH (Millipore-Sigma G8795, 1:5000) antibodies. Secondary IR antibodies were purchased from Li-COR.

### PARP14 cellular ADP-ribosylation assays

293T cells were transfected with GFP-PARP14 WT or GFP-PARP14 G832E plasmid using the jetOPTIMUS transfection reagent with 0.4 µg DNA/well and with a ratio of 1:1 µL transfection reagent: µg DNA. The media was exchanged 4 hours post-transfection. The next day, cells were dosed with RBN012759 for 4 hours. Cells were washed once with cold PBS and the plate was frozen at -80°C prior to lysis. Samples were thawed on ice, lysed with a cytosolic lysis buffer [50 mM HEPES pH 7.4, 150 mM NaCl, 1 mM MgCl_2_, 1% tritonX-100 supplemented with fresh 1 mM TCEP, 1X cOmplete EDTA-free Protease Inhibitor Cocktail (Roche) and 30 µM Phthal01 (panPARP inhibitor) [46]], and clarified by centrifugation at 14,000 rpm at 4°C for 10 minutes. The supernatant containing total protein was quantified using the Bradford assay reagent (Bio-Rad). Lysates were normalized and sample buffer was added to 1X (10% glycerol, 50mM Tris-Cl (pH 6.8), 2% SDS, 1% β-mercaptoethanol, 0.02% bromophenol blue). Samples were boiled at 95°C for 5 min and resolved in 4-20% Mini-PROTEAN® TGX™ Precast Protein Gels (Bio-Rad). Proteins were transferred to a nitrocellulose membrane and probed overnight at 4°C with primary antibodies, followed by incubation with goat anti-rabbit (1:10000, Jackson Immuno Reseach Labs) or goat anti-mouse (1:5000, Invitrogen) HRP-conjugated secondary antibodies. The blots were then incubated SuperSignal™ West Pico substrate (Thermo Scientific) and imaged on a ChemiDoc Gel Imaging System (Bio-Rad). Primary antibodies used for this experiment were: Poly/Mono-ADP ribose antibody (Cell Signaling Technology, E6F6A, 1:2000), Tubulin (Cell Signaling Technology, DM1A, 1:2000), GFP (Proteintech, pabg1, 1:1000), Mono-ADP-Ribose antibody 33204 (Bio-Rad, HCA354, 2 µg/ml), and PARP14 (Millipore Sigma, HPA012063, 1:1000).

### Real-Time (RT-qPCR) Analysis

RNA was isolated from respective cells using Trizol (Invitrogen). Briefly, cells were collected at indicated time points in Trizol (Invitrogen) and RNA was isolated as per manufacturer’s instructions. cDNA was prepared using M-MuLV-reverse transcriptase per the manufacturer’s instructions (NEB). qPCR was performed using PowerUp SYBR green master mix (Applied Biosystems) and primers listed in **Table S3**. Cycle threshold values were normalized to hypoxanthine phosphoribosyltransferase (HPRT), glyceraldehyde-3-phosphate dehydrogenase (GAPDH), or 18S rRNA levels using the ΔCt method.

### RNAseq

RNA was isolated from infected BMDMs with and without treatment, isolated from C57BL6 mice as described above. Library preparation with indexing was performed by the University of Kansas Genome Sequencing core facility with NEB Next RNA Library kit (NEB). RNAseq was performed using an Illumina NextSeq2000 high-output system with paired-end reads of 50 bp. DESeq2 was used to identify DEGs between the BMDM without treatment and MHV-JHM N1347A infected and BMDM with 100nM PARP14 inhibitor treatment and N1347A infected samples using simply “treatment” as a factor. DEGs were identified based on the false-discovery rate corrected *P*-value (P_ADJ_) and log_2_-fold-change of (log_2_FC) between the samples. Genes were considered up-regulated in an MHV-JHM-infected sample if P_ADJ_ < 0.05 and log_2_FC > 0.6, which is nearly equivalent to a 1.5-fold increase. Similarly, genes were considered down-regulated if P_ADJ_ < 0.05 and log_2_FC < −0.6, or a 1.5-fold decrease. DEGs were subjected to gene ontology analysis using the Database for Annotation, Visualization and Integrated Discovery (DAVID: https://david.ncifcrf.gov/).

### Primers

Primers were used at 10μM concentration and the primers sequences for human HPRT, IFNβ, GAPDH [47], CXCL10 and mouse HPRT, IFNβ and ISG15 have been previously published [25]. RNAseq data was corroborated by qPCR using primer sequences CASP1F TACACGTCTTGCCCTCATTATC, CASP1R CTCCAGCAGCAACTTCATTTC, FOXO3F TTTCCTACGTCTCCTGCTAATG, FOXO3R CTCCTTAGGCTGACGCTAATC, RNF111F CGACTTCATCACCTCCAGTTAG, RNF111R GCTCCATCCAATCCTGAAGAA. Dentritic cell verification was done using qPCR using the primers CXCR1F TTCCCATCTGCTCAGGACCTC and CXCR1R ATTTCCCACCAGACCGAACG.

### Immunofluorescence Analysis and Imaging

Immunofluorescence to detect NiV was performed as described as follows. Briefly, formalin-fixed cells were washed 4 times with phosphate-buffered saline (PBS), permeabilized with a mixture of acetone and methanol (1:1, *v*/*v*) for 5 min at −20 °C, washed 4 times with PBS, treated with 0.1 M glycine for 5 min at room temperature, washed 4 times with PBS, and then incubated in blocking solution (2% bovine serum albumin, 0.2% Tween 20, 3% glycerol, and 0.05% sodium azide in PBS) for 1 h at room temperature. The primary antibody used for the detection of NiV (mouse hyperimmune ascitic fluid diluted 1:500 in blocking solution, BEI NR-48961) was added and incubated overnight at 4°C. Cells were washed 4 times with PBS and then incubated with secondary antibody (goat α-mouse AlexaFluor 488 diluted 1:100 in blocking solution, Invitrogen) in the presence of 4′,6-diamidino-2-phenylindole (DAPI; 200 ng/mL). rEBOV-ZsG-infected cells were stained with DAPI (200 ng/mL). Infections and immunofluorescence analyses were performed three times in independent experiments.

Samples were imaged at 4x magnification on a Nikon Ti2 Eclipse inverted fluorescent microscope and Photometrics Prime BSI camera using NIS Elements AR software. One representative image from each well, covering an area of 10.87 mm^2^, was collected for quantification.

### Statistics

A Student’s *t* test or one-way ANOVA were used to analyze differences in mean values between groups. All results are expressed as means ± standard errors of the means (SEM). P values of ≤0.05 were considered statistically significant (*, P≤0.05; **, P≤0.01; ***, P≤0.001; ****, P ≤0.0001; n.s., not significant). “ns” stands for data with P value >0.05 and hence is not significant. Fold changes are designated by a number followed by X above the significance values.

### Data and materials availability

All the RNAseq reads data will be deposited in NCBI (Bioproject ID: PRJNA1139810; Biosample ID: SAMN42785804 (DMSO) and SAMN42786772(PARP14 inhibitor treatment)) and will be made public upon acceptance. All other data will be available through FigShare at the time of acceptance. 10.6084/m9.figshare.c.7766243

## ACKNOWLEDGMENTS

We thank members of the Davido and Orozco labs for valuable discussion. We thank Stanley Perlman, Susan Weiss, Mario Niepel, Asit Pattnaik, and Heinz Feldmann for reagents. Bioinformatic consultation was provided by the KU Center for Genomics. Research reported in this publication was made possible in part by the services of the KU Genome Sequencing Core which is supported by NIH/NIGMS under grant award number P30GM145499, and the KU Flow Cytometry Core, which is supported in part by the NIH/NIGMS under grant award number P20GM113117. Transgenic mice were generated at the University of Iowa Genome Editing Core Facility directed by William Paradee, PhD and supported in part by grants from the NIH and from the Roy J. and Lucille A. Carver College of Medicine. We wish to thank Norma Sinclair, Patricia Yarolem, Joanne Schwarting and Rongbin Guan for their technical expertise in generating transgenic mice. We also thank Mitchell White (Boston University) for excellent technical assistance.

## Author contributions

Conceptualization: SP, ARF

Methodology: SP, ELS, DSB, NS, MA, RCO, CSS, MSC, DJD, AJH, ARF

Investigation: SP, PS, HH, AF, JJP, ELS, DSB, YC, NS, RCO, CSS, DJD, AJH, ARF

Visualization: SP, PS, HH, AF, JJP, ELS, DSB, YC, RCO, CSS, MSC, DJD, AJH, ARF

Supervision: SP, EM, RCO, CSS, MSC, DJD, AJH, ARF

Writing—original draft: SP, ARF

Writing—review & editing: SP, PS, HH, AF, JJP, ELS, DSB, YC, NS, JA, EM, RCO, CSS, MSC, DJD, AJH, ARF

## SUPPLEMENTAL FIGURE LEGENDS

**Fig. S1.** A) WT and PARP14KO clone 23 (C23) A549 cells were transfected with 0.5μg/mL poly(I:C), at 18 hpt RNA was isolated from cells and the level of IFNβ mRNA was quantified using qPCR using ΔCt method. Data shown in D are from 1 experiment and are representative of 3 independent experiments with N=3 for each experiment. B) WT and PARP14 KO NHDF cells were transfected with 0.5μg/mL poly(I:C), at 18 hpt RNA was isolated from cells and the level of IFNβ mRNA was quantified using qPCR using ΔCt method. Data shown in A are from 1 experiment and are representative of 3 independent experiments with N=3 for each experiment. C) 293T cells were mock transfected (M) or transfected with plasmid expressing GFP-PARP14 G832E. At 24 hpt, cells were treated with PARPi for 4 hrs. Cells were then collected and analyzed by immunoblot with indicated antibodies. Data shown in B are from 1 experiment representative of 2 independent experiments. D) A549 WT cells were treated with either DMSO, negative control (CTL) or PARP14 Degron (DEG) for 6 hrs and then wither mock transfected or transfected with 0.5μg/mL of poly(I:C) with the corresponding treatments. At 18 hpt cell lysates were collected and PARP14 protein levels were determined by immunoblotting using β-actin as a loading control. E) WT A549 cells transfected with poly(I:C) were treated with 1 μM PARP14 Degron control (CTL) or PARP14 Degron (DEG) for 18 hours. RNA was isolated from cells at 18 hpt and IFNβ mRNA was quantified by qPCR using ΔCt method. Data shown in E are from 1 experiment and are representative of 3 independent experiments with N=3 for each experiment.

**Fig. S2.** A) Floxed Parp14 *cre*-(*Parp14+/+*) and *cre*+ (*Parp14-/-*) BMDMs were treated with 4-OHT and 24 hours later were infected with MHV WT and N1347A at an MOI of 0.1. At 12hpi, cell lysates were harvested and levels of PARP14 and β-actin were determined by immunoblotting. B) *Parp14+/+* and *Parp14-/-* BMDMs were infected with WT and N1347A MHV-JHM at an MOI of 0.1. RNA was collected 12 hpi and ISG15 mRNA levels were quantified using qPCR using ΔCt method. Data shown in B are from 1 experiment and are representative of 3 independent experiments with N=3 biological replicates for each experiment. C) RNA was isolated from BMDMs and DCs and CXCR1 mRNA levels were quantified using qPCR using ΔCt method. Data shown in C are the combined data from 2 independent experiments with N=6 biological replicates for each group.

**Fig. S3. PARP14 does not impact LCMV infection.** A) Gating strategy used to identify LCMV^cl13^(+) cells by flow cytometry. B) WT and PARP14 KO BMDMs were infected with LCMV^cl13^ at an MOI of 1 and the % of LCMV^cl13^ (+) cells were quantified by flow cytometry. These data are from 1 experiment representative of 3 independent experiments. N=3 per group.

**Fig S4. PARP14 knockout (KO) in cell lines using CRISPR/Cas9.** Using CRISPR-Cas9 based methods, we developed A549-ACE PARP14 KO cells. Western blot confirmed that PARP14 was absent in multiple clones of A549-ACE2 lung epithelial cells.

**Fig S5. Reduced VSV replication in A549 PARP14 KO cells is not due to IFN.** A) A549 WT and PARP14 KO cells were infected with VSV-GFP at an MOI of 1 and treated with 1 μM JAK inhibitor (JAKi). RNA was isolated from cells at 10 hpi and IFN-*β* mRNA was quantified by qPCR using ΔCt method. Data shown in A are from 1 experiment and are representative of 2 independent experiments with N=3 biological replicates for each experiment. B) A549 WT and PARP14 KO cells were infected with VSV-GFP at an MOI of 1 and treated with 1 μM JAK inhibitor (JAKi). RNA was isolated from cells at 10 hpi and PARP14 mRNA was quantified by qPCR using ΔCt method. Data shown in B are from 1 experiment and are representative of 2 independent experiments with N=3 biological replicates for each experiment. C) A549 WT and PARP14 KO cells were infected with VSV-GFP at an MOI of 1 and treated with 1 μM JAK inhibitor (JAKi). At 10 hpi, cells and supernatants were collected, and the progeny virus was quantified by TCID_50_. Results in C are from one experiment representative of three independent experiments with N=3 biological replicates for each experiment.

