## Supplemental Figures for "PARP14 is an interferon (IFN)-induced host factor that promotes IFN production and affects the replication of multiple viruses"

### Slide 1
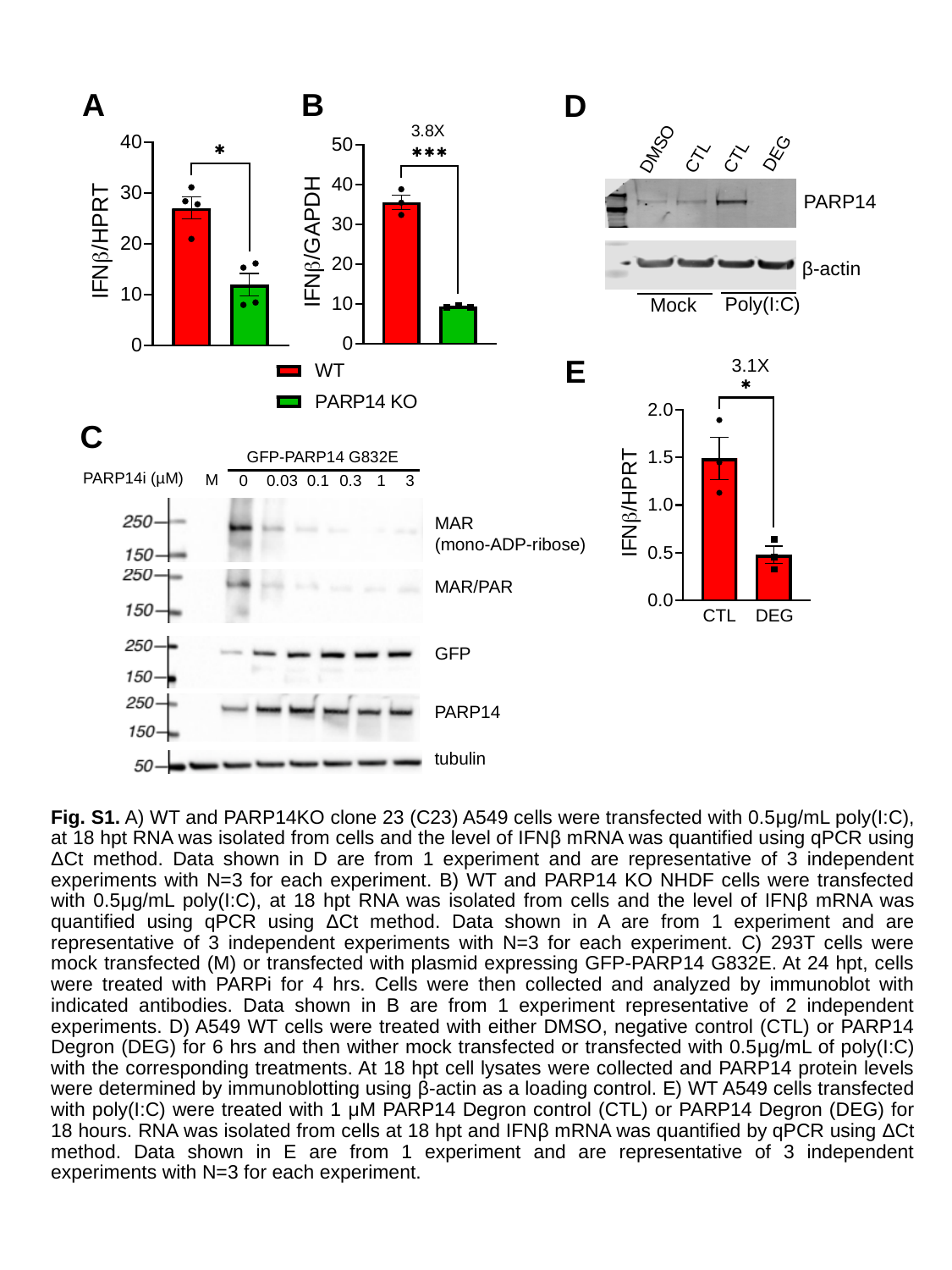

A
B
D
DMSO
CTL
DEG
CTL
PARP14
β-actin
Poly(I:C)
Mock
E
C
GFP-PARP14 G832E
PARP14i (µM)
M
0
0.03
0.1
0.3
1
3
MAR
(mono-ADP-ribose)
MAR/PAR
CTL
DEG
GFP
PARP14
tubulin
Fig. S1. A) WT and PARP14KO clone 23 (C23) A549 cells were transfected with 0.5μg/mL poly(I:C), at 18 hpt RNA was isolated from cells and the level of IFNβ mRNA was quantified using qPCR using ΔCt method. Data shown in D are from 1 experiment and are representative of 3 independent experiments with N=3 for each experiment. B) WT and PARP14 KO NHDF cells were transfected with 0.5μg/mL poly(I:C), at 18 hpt RNA was isolated from cells and the level of IFNβ mRNA was quantified using qPCR using ΔCt method. Data shown in A are from 1 experiment and are representative of 3 independent experiments with N=3 for each experiment. C) 293T cells were mock transfected (M) or transfected with plasmid expressing GFP-PARP14 G832E. At 24 hpt, cells were treated with PARPi for 4 hrs. Cells were then collected and analyzed by immunoblot with indicated antibodies. Data shown in B are from 1 experiment representative of 2 independent experiments. D) A549 WT cells were treated with either DMSO, negative control (CTL) or PARP14 Degron (DEG) for 6 hrs and then wither mock transfected or transfected with 0.5μg/mL of poly(I:C) with the corresponding treatments. At 18 hpt cell lysates were collected and PARP14 protein levels were determined by immunoblotting using β-actin as a loading control. E) WT A549 cells transfected with poly(I:C) were treated with 1 μM PARP14 Degron control (CTL) or PARP14 Degron (DEG) for 18 hours. RNA was isolated from cells at 18 hpt and IFNβ mRNA was quantified by qPCR using ΔCt method. Data shown in E are from 1 experiment and are representative of 3 independent experiments with N=3 for each experiment.

### Slide 2
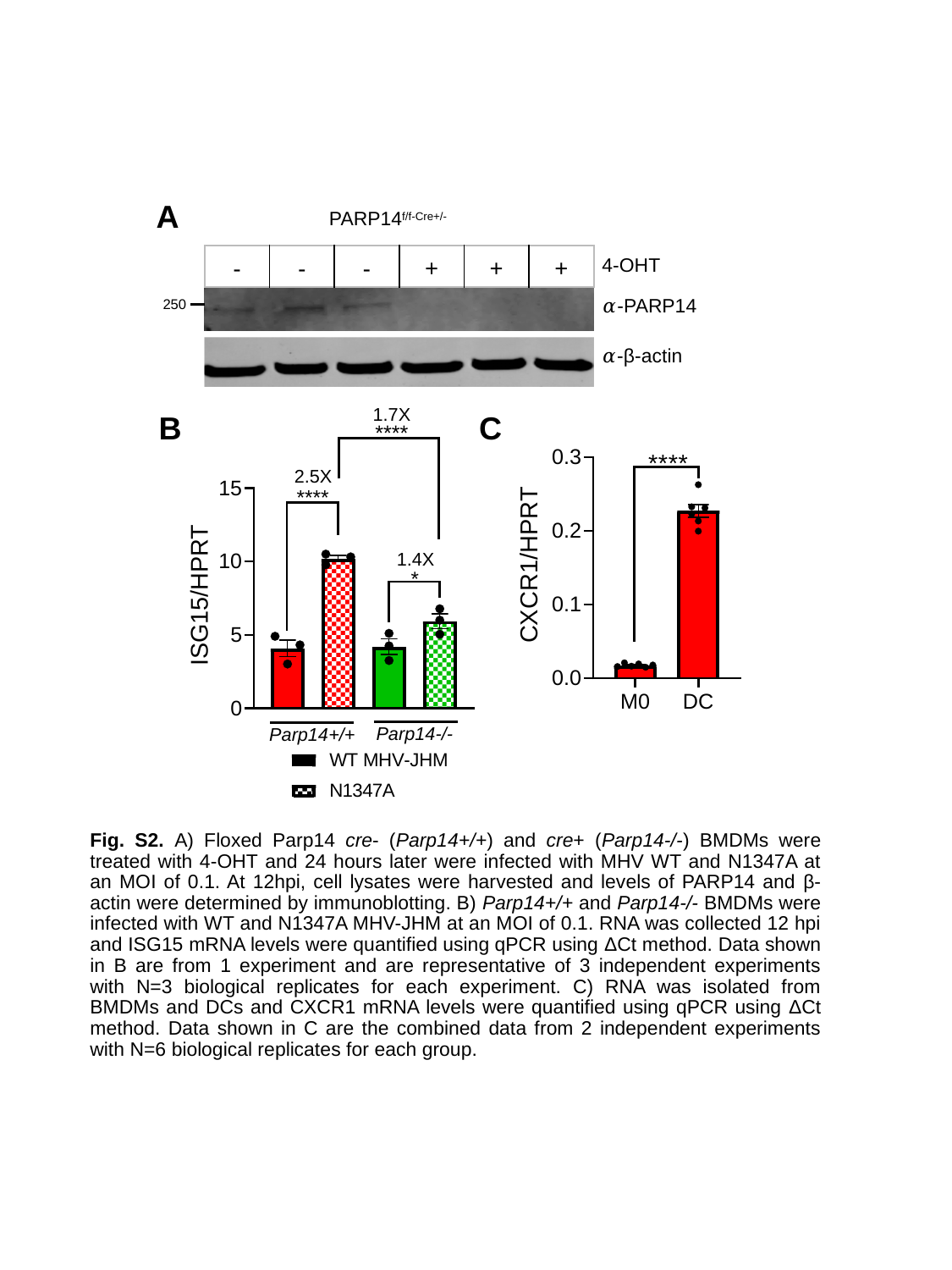

A
PARP14^f/f-Cre+/-^
| - | - | - | + | + | + |
| --- | --- | --- | --- | --- | --- |
4-OHT
𝛼-PARP14
250
𝛼-β-actin
C
B
Fig. S2. A) Floxed Parp14 cre- (Parp14+/+) and cre+ (Parp14-/-) BMDMs were treated with 4-OHT and 24 hours later were infected with MHV WT and N1347A at an MOI of 0.1. At 12hpi, cell lysates were harvested and levels of PARP14 and β-actin were determined by immunoblotting. B) Parp14+/+ and Parp14-/- BMDMs were infected with WT and N1347A MHV-JHM at an MOI of 0.1. RNA was collected 12 hpi and ISG15 mRNA levels were quantified using qPCR using ΔCt method. Data shown in B are from 1 experiment and are representative of 3 independent experiments with N=3 biological replicates for each experiment. C) RNA was isolated from BMDMs and DCs and CXCR1 mRNA levels were quantified using qPCR using ΔCt method. Data shown in C are the combined data from 2 independent experiments with N=6 biological replicates for each group.

### Slide 3
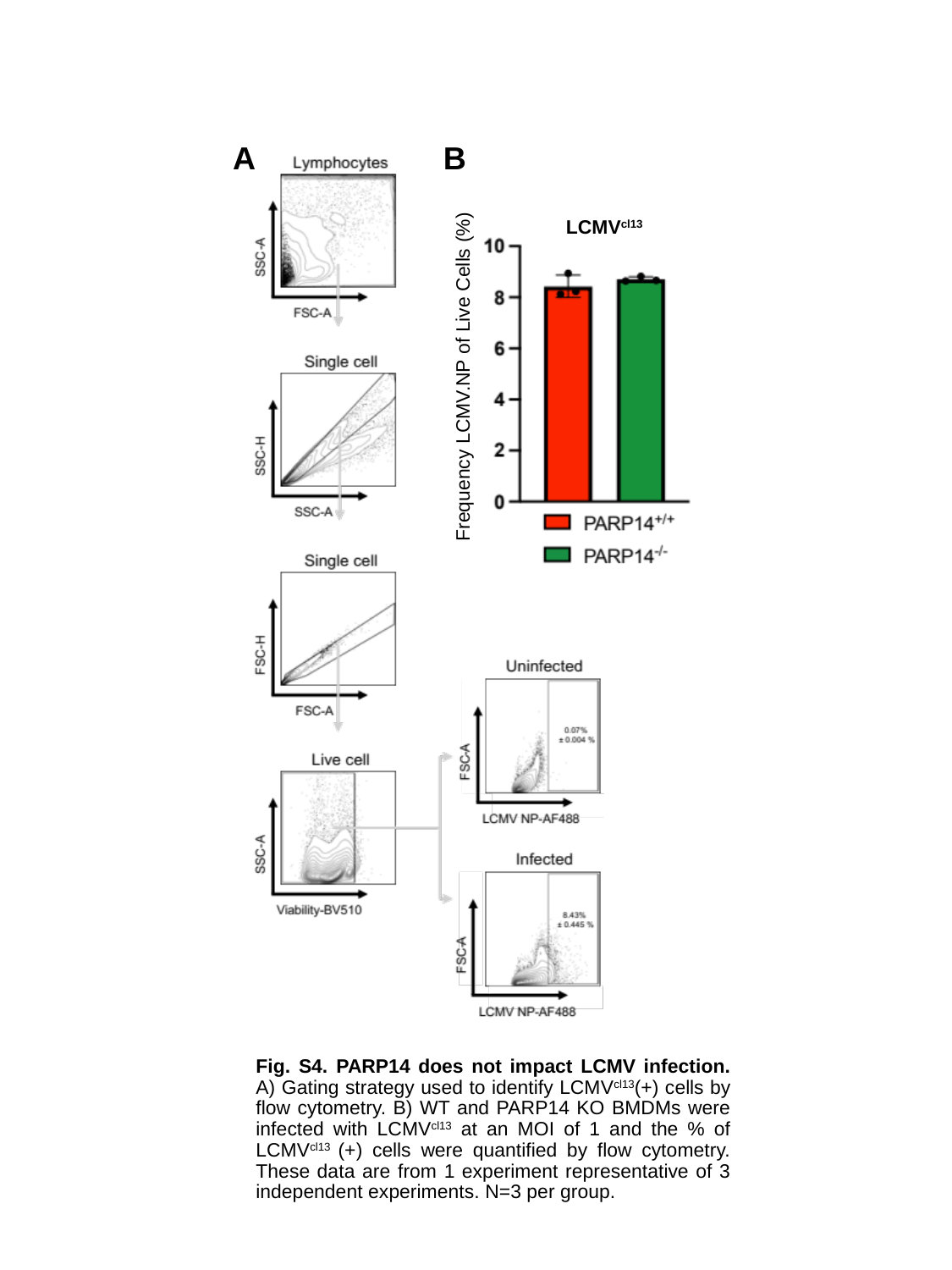

B
A
LCMV^cl13^
Frequency LCMV.NP of Live Cells (%)
Fig. S4. PARP14 does not impact LCMV infection. A) Gating strategy used to identify LCMV^cl13^(+) cells by flow cytometry. B) WT and PARP14 KO BMDMs were infected with LCMV^cl13^ at an MOI of 1 and the % of LCMV^cl13^ (+) cells were quantified by flow cytometry. These data are from 1 experiment representative of 3 independent experiments. N=3 per group.

### Slide 4
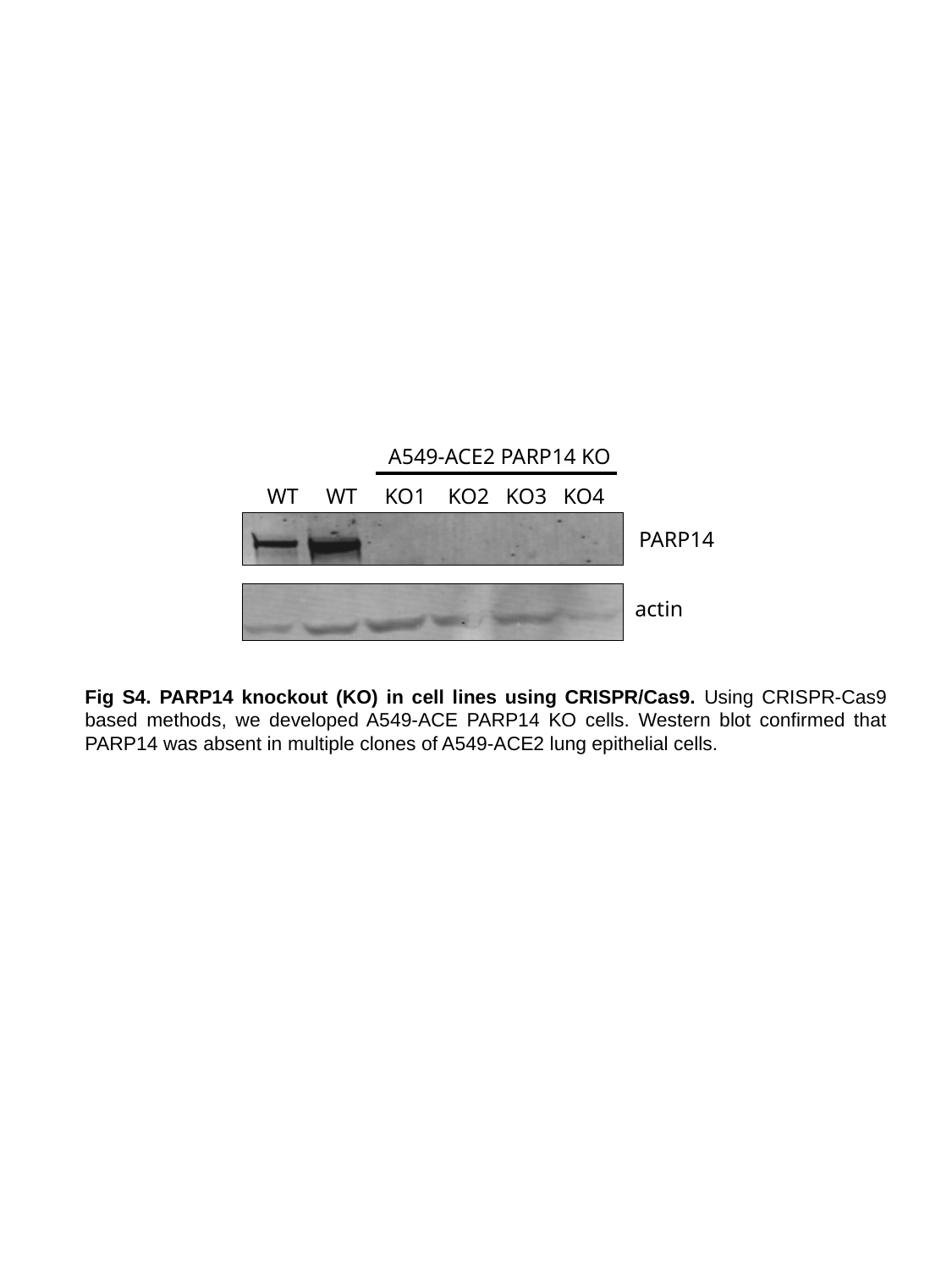

A549-ACE2 PARP14 KO
 WT WT KO1 KO2 KO3 KO4
PARP14
actin
Fig S4. PARP14 knockout (KO) in cell lines using CRISPR/Cas9. Using CRISPR-Cas9 based methods, we developed A549-ACE PARP14 KO cells. Western blot confirmed that PARP14 was absent in multiple clones of A549-ACE2 lung epithelial cells.

### Slide 5
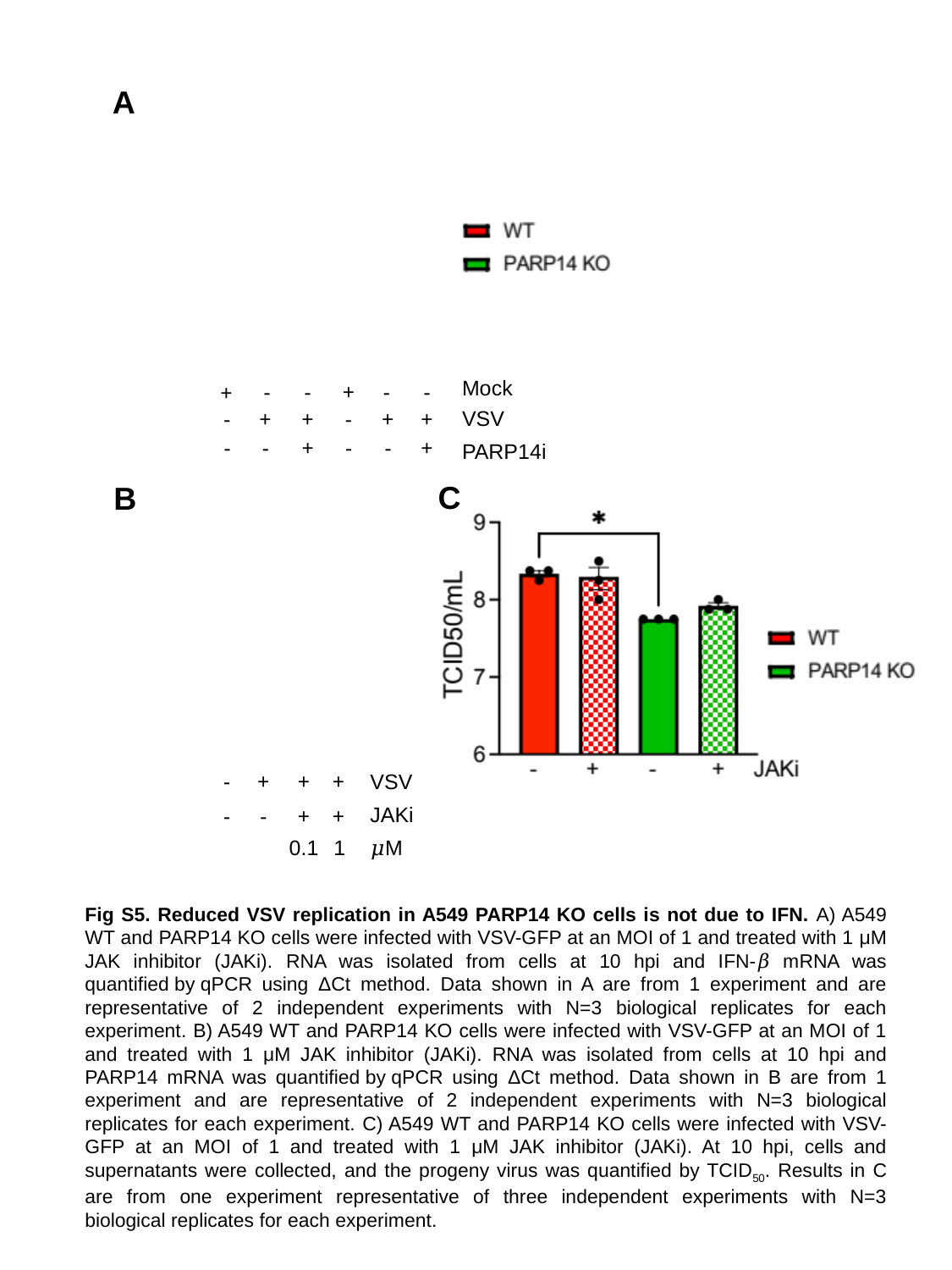

A
Mock
+
-
-
-
-
+
VSV
-
+
+
-
+
+
-
-
+
-
-
+
PARP14i
C
B
-
+
+
+
VSV
JAKi
-
-
+
+
1
0.1
𝜇M
Fig S5. Reduced VSV replication in A549 PARP14 KO cells is not due to IFN. A) A549 WT and PARP14 KO cells were infected with VSV-GFP at an MOI of 1 and treated with 1 μM JAK inhibitor (JAKi). RNA was isolated from cells at 10 hpi and IFN-𝛽 mRNA was quantified by qPCR using ΔCt method. Data shown in A are from 1 experiment and are representative of 2 independent experiments with N=3 biological replicates for each experiment. B) A549 WT and PARP14 KO cells were infected with VSV-GFP at an MOI of 1 and treated with 1 μM JAK inhibitor (JAKi). RNA was isolated from cells at 10 hpi and PARP14 mRNA was quantified by qPCR using ΔCt method. Data shown in B are from 1 experiment and are representative of 2 independent experiments with N=3 biological replicates for each experiment. C) A549 WT and PARP14 KO cells were infected with VSV-GFP at an MOI of 1 and treated with 1 μM JAK inhibitor (JAKi). At 10 hpi, cells and supernatants were collected, and the progeny virus was quantified by TCID_50_. Results in C are from one experiment representative of three independent experiments with N=3 biological replicates for each experiment.
